# Cell specific single viral vector CRISPR/Cas9 editing and genetically encoded tool delivery in the central and peripheral nervous systems

**DOI:** 10.1101/2023.10.10.561249

**Authors:** Jamie C. Moffa, India N. Bland, Jessica R. Tooley, Vani Kalyanaraman, Monique Heitmeier, Meaghan C. Creed, Bryan A. Copits

## Abstract

Gene manipulation strategies using germline knockout, conditional knockout, and more recently CRISPR/Cas9 are crucial tools for advancing our understanding of the nervous system. However, traditional gene knockout approaches can be costly and time consuming, may lack cell-type specificity, and can induce germline recombination. Viral gene editing presents and an exciting alternative to more rapidly study genes of unknown function; however, current strategies to also manipulate or visualize edited cells are challenging due to the large size of Cas9 proteins and the limited packaging capacity of adeno-associated viruses (AAVs). To overcome these constraints, we have developed an alternative gene editing strategy using a single AAV vector and mouse lines that express Cre-dependent Cas9 to achieve efficient cell-type specific editing across the nervous system. Expressing Cre-dependent Cas9 in specific cell types in transgenic mouse lines affords more space to package guide RNAs for gene editing together with Cre-dependent, genetically encoded tools to manipulate, map, or monitor neurons using a single virus.

We validated this strategy with three commonly used tools in neuroscience: ChRonos, a channelrhodopsin, for manipulating synaptic transmission using optogenetics; GCaMP8f for recording Ca2+ transients using fiber photometry, and mCherry for anatomical tracing of axonal projections. We tested these tools in multiple brain regions and cell types, including GABAergic neurons in the nucleus accumbens (NAc), glutamatergic neurons projecting from the ventral pallidum (VP) to the lateral habenula (LHb), dopaminergic neurons in the ventral tegmental area (VTA), and parvalbumin (PV)-positive proprioceptive neurons in the periphery. This flexible approach should be useful to identify novel genes that affect synaptic transmission, circuit activity, or morphology with a single viral injection.

## Introduction

Advances in genetic targeting strategies have been instrumental for understanding how specific genes influence normal and pathological functions in the nervous system. Furthermore, combining these manipulations with genetically encoded fluorophores, calcium indicators, and optogenetic or chemogenetic tools provides powerful approaches to understand how specific genes regulate synaptic, circuit, and behavioral phenotypes. Fluorescent proteins permit direct visualization of axonal projections, dendritic arbors, and the synapses connecting them (Chen et al., 2017; Meltzer et al., 2023; Tashiro et al., 2006; Young et al., 2008; J.-H. Zhang et al., 2011; X. Zhang et al., 2022), while calcium sensors such as GCaMP enable optical recording of cellular activity *in vivo* (Brandner et al., 2023; McQuillan et al., 2022; Zhou et al., 2022). Opsins, like channelrhodopsin (ChR), or chemogenetic tools such as designer receptors exclusively activated by designer drugs (DREADDs), can directly manipulate the activity of defined cell types to understand how various genes regulate their function (Garrido et al., 2022; Hadjas et al., 2020; Rapanelli et al., 2017; Stuber et al., 2010).

Global gene knockout (KO) mice have revolutionized our understanding of foundational concepts in neuroscience. However, global knockouts have significant limitations: some are lethal (Gangloff et al., 2004; E. Li et al., 1992), and all global knockouts lack temporal, spatial, and cell-type specificity. The Cre-Lox system offers significant advantages to manipulate gene expression with increased specificity (Orban et al., 1992). This can involve transgenic crosses to improve cellular specificity (Tsien et al., 1996) or viral strategies that enable increased spatial and temporal control of gene deletion (Kaspar et al., 2002). However, generating cKO lines have several significant drawbacks. First, Cre-driver promoters can become transiently activated during development or have leaky expression, leading to off-target expression of Cre and gene deletion in unintended cell types (Lin Luo et al., 2020; Madisen et al., 2010; Song & Palmiter, 2018). Second, this approach necessitates multiple crosses, thus sacrificing time while increasing colony sizes and maintenance costs. These limitations are amplified when studying more than one gene. Third, expression of genetically encoded tools in cKO neurons may require introduction of yet an additional mouse line expressing a Cre-dependent reporter gene, further complicating breeding strategies.

Viral strategies can overcome many of the drawbacks faced by generating cKO lines while retaining the benefits of highly efficient gene deletion (Kaspar et al., 2002). This approach requires only a floxed mouse line and injection of Cre-expressing viruses, thus enabling more precise control over the timing and location of gene knockouts. While viral vector promoters can restrict expression to specific cell types, well-characterized promoters that drive sufficient, specific expression are limited to a few cell types (Sjulson et al., 2016). Many viral promoters can also be leaky (Sjulson et al., 2016), and are limited by viral vector packaging size (Dong et al., 1996). Both cKO strategies require a new floxed line for each target gene, making it cost-prohibitive to study genes of unknown function or to screen multiple genes of interest. In the era of rapidly developing large-scale transcriptomic and genome-wide association studies (GWAS), a higher-throughput tool is needed to study multiple candidate genes without creating a new mouse line for each gene of interest.

CRISPR/Cas9 is a powerful tool that addresses many of the drawbacks of the Cre-Lox system and has opened new horizons for studying gene function. It takes advantage of bacterial Cas9 proteins, which complex with short segments of guide RNA (gRNA) to target specific regions of DNA and cause a double-stranded break (Jiang & Doudna, 2017; Jinek et al., 2012; Kalamakis & Platt, 2023). Cellular repair of these breaks via non-homologous end joining (NHEJ) often results in frame-shifting insertions or deletions generating a premature stop codon. While CRISPR-Cas9 gene editing in the nervous system was slower to develop, successful editing has recently been demonstrated for multiple genes across diverse brain regions and cell types using viral vectors containing a targeting gRNA and Cre-dependent Cas9 injected into transgenic Cre-driver mice (Castro et al., 2021; Fellinger et al., 2021; Gunduz-Cinar et al., 2023; Hunker et al., 2020; H. Li et al., 2022; Liu et al., 2022; Soden et al., 2023; Swiech et al., 2015). This system allows for cell-type specificity and precise spatial and temporal gene deletion to study neuronal sub-populations in specific brain regions. As gene editing is achieved via gRNA targeting rather than floxed mouse lines, multiple candidate genes can be screened far more quickly and at lower costs. However, current iterations of this strategy require injection of two AAV vectors to achieve both editing and transgene expression. This is because Cas enzymes are large; only a few are small enough to be packaged into AAVs, which have a maximum capacity of ∼4.7 kb (Challis et al., 2019; Dong et al., 1996; Grieger & Samulski, 2005; Hunker et al., 2020) Thus, a second viral vector carrying a transgene for manipulating, imaging, or tracing edited neurons must be co-injected. As transfection efficiency for any single virus is less than 100%, introducing multiple viruses compounds this inefficiency, resulting in some edited neurons that are not labeled with the transgene, and some unedited neurons that nevertheless contain the transgene. This imprecision could lead to data misinterpretations from labeled, but unedited neurons.

We designed and validated a novel approach that addresses these limitations with two-vector CRISPR/Cas9 editing techniques. We first modified existing viral vectors with simple cloning sites for gRNA insertion to co-express Cre-dependent transgenes for visualization, manipulation, and recording of target neurons. This enables Cas9-mediated editing and transgene expression only in specific labeled neuronal populations. As this method only uses a single viral vector, this ensures that the edited population of neurons fully overlaps with transgene expression. To circumvent the size limitation of packaging Cas9 in an AAV vector, we crossed Cre-dependent Cas9 mouse lines (Chiou et al., 2015; Platt et al., 2014) to Cre-driver lines to enable cell-type specific Cas9 expression. We validated this approach with three commonly used tools in neuroscience: ChRonos, a channelrhodopsin, for manipulating synaptic transmission using optogenetics; GCaMP8f for recording Ca2+ transients using fiber photometry, and mCherry for anatomical tracing of axons. This new flexible approach can be used to screen multiple target genes for effects on synaptic transmission, circuit activity, or morphology with a single viral injection.

## Results

### Combining CRISPR/Cas9 with optogenetics to study synaptic physiology

To test the utility of our single-vector approach for transgene expression and CRISPR/Cas9 editing, we used optogenetic slice recordings while targeting proteins regulating neurotransmitter release. We focused on the vesicular GABA transporter (*Slc32a1* or *Vgat*) that packages GABA into synaptic vesicles, where effective editing should reduce the amplitude of optogenetically-evoked inhibitory postsynaptic currents (oIPSCs). To do this, we first cloned the Cre-dependent fast channelrhodopsin ChRonos-GFP (Klapoetke et al., 2014) into an empty pX552 gRNA viral vector (Swiech et al., 2015) (**Figure 1A**). We next used the bioinformatics database CRISPOR (Concordet & Haeussler, 2018) to design a gRNA against *Vgat* and a nearly-identical control gRNA that differed from the active construct by three thymidine residues near the protoadjacent spacer motif (PAM) (Hunker et al., 2020) (**Figure 1B**). We inserted either the active or control gRNA into the *SapI* cloning site of the pX552:U6:gRNA:Ef1a:FLEx:ChRonos-GFP construct, and packaged these into AAV5 viruses to efficiently target CNS neurons. To selectively express Cas9 in GABAergic neurons, we crossed *Vgat*-Cre mice (Vong et al., 2011) to a Cre-dependent Cas9 line (Platt et al., 2014). We then injected 6-8 week old adults from the F1 generation with control or active virus in the nucleus accumbens (NAc) (**Figure 1C**). We chose the NAc because it contains a high density of locally-projecting GABAergic neurons to test the efficacy of VGAT manipulations.

**Figure 1.**
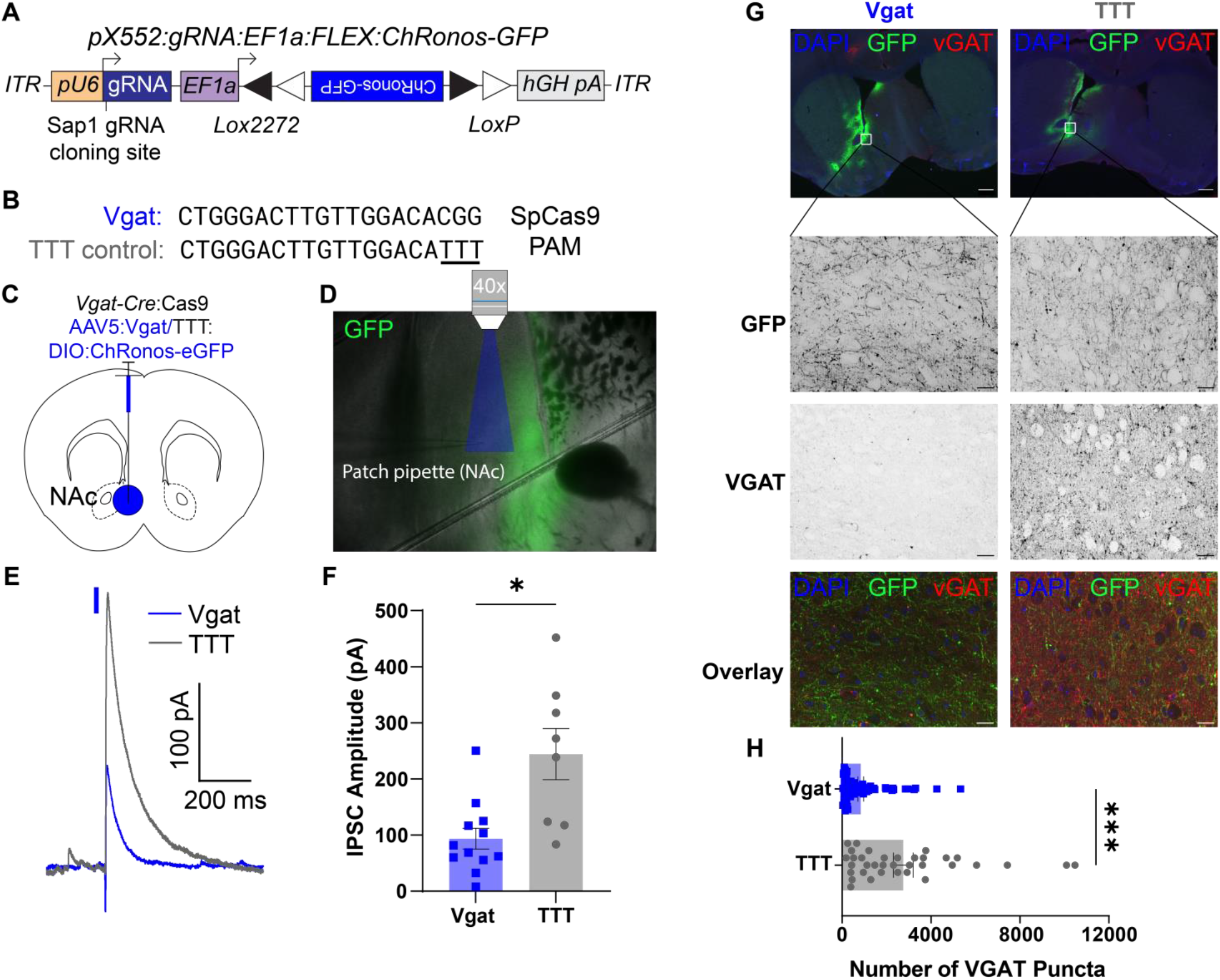
ChRonos-GFP coupled with *Vgat* editing demonstrates reduced inhibitory synapse function. (**A**) Schematic of pX552 viral vector with Cre-dependent ChRonos-GFP transgene and Vgat gRNA for gene editing. (**B**) Sequences of gRNA targeting *Vgat* (top) and modified TTT control (bottom). (**C**) Diagram of virus injection into the NAc of 6-8 week *Vgat*-Cre:Cas9 mice. (**D**) Representative image of slice electrophysiology setup, 4x magnification. Area of blue light stimulation of ChRonos shown in blue. ChRonos-GFP expression shown in green. (**E**) Representative oIPSC traces from *Vgat-*edited (blue) and TTT control (grey) neurons. Traces aligned to light pulse onset (blue rectangle). The representative edited IPSC trace is an average of 5 consecutive sweeps; the representative control IPSC trace is an average of 4 consecutive sweeps. (**F**) Quantification of evoked IPSC amplitude. Each point represents the average peak inhibitory current amplitude after blue light stimulus from 3-10 consecutive sweeps. The average Vgat IPSC amplitude was 93.56±18.58 pA (n=12 cells from 3 animals), and the average TTT IPSC amplitude was 244.3±45.64 pA (n=8 cells from 2 animals) (p=0.013*). (**G**) Representative VGAT IHC images for Vgat-edited (left) and TTT (right) conditions. **Top:** 10x images showing GFP expression for virus injection distribution (green), VGAT puncta labeling (red) and DAPI counterstain (blue). **2^nd^ Row:** 60x images showing GFP expression in the NAc of Vgat (left) and TTT (right) slices. **3^rd^ Row:** 60x images showing VGAT labeling in Vgat-edited (left) and TTT (right) slices. **Bottom:** Overlay of GFP (green), VGAT (red), and DAPI (blue). 10x scale bars=100 µm; 60x scale bars=20 µm. (**H**) Quantification of number of VGAT puncta in Vgat-edited (top) or TTT (bottom) sections. Each point represents the number of VGAT puncta in one 241.56 x 181.17 µm NAc section imaged at 60x. The average number of puncta in Vgat-edited sections was 838.9±128.3 puncta (n=35 sections from 4 animals); the average number of puncta in TTT sections was 2751±442.2 puncta (n=70 sections from 6 animals) (p=0.0002***). All comparisons were done using a 2-tailed, unpaired t-test with Welch’s correction for unequal variances. All data are reported as mean±SEM.

After 6 weeks, we performed whole-cell patch-clamp recordings from acute slices of the NAc. We recorded pharmacologically-isolated oIPSCs at 0 mV by stimulating ChRonos with blue light (1 ms, 1 mW/mm2) (**Figure 1D**). We found that oIPSC amplitudes in neurons from mice injected with active virus were significantly reduced compared to control (**Figure 1E,F**). To validate that our viral vector reduces VGAT expression in GABAergic NAc neurons, we injected either the active or control virus into the nucleus accumbens (NAc) of adult *Vgat*-Cre/lsl-Cas9 mice, as above. After 6 weeks to allow for transgene expression and editing, we analyzed VGAT puncta using immunohistochemistry (IHC) and found that mice injected with active gRNA had significantly fewer VGAT puncta in GFP+ regions of the NAc, compared with control mice (**Figure 1G,H**). These data demonstrate that our single-vector approach can be used to edit VGAT, visualize edited neurons, and investigate functional changes in edited neurons using optogenetics.

While our results targeting GABAergic neurons in the NAc suggest efficient editing, it is possible that the variation in oIPSC amplitudes could be due to differences in viral injection placement or efficiency, leading to optogenetic activation of different numbers of presynaptic terminals between the two groups. To rule out this possibility, we next targeted *Vglut2*+ neurons that project from the ventral pallidum (VP) to the lateral habenula (LHb) and co-release both glutamate and GABA (Faget et al., 2018; Tooley et al., 2018). Since glutamate packaging into synaptic vesicles requires vesicular glutamate transporters (VGLUTs), synaptic glutamate release should not be affected by *Vgat* editing. Using this internal control, we can then normalize oIPSC amplitudes to optogenetically-evoked excitatory postsynaptic current (oEPSC) amplitudes to control for the number of terminals activated by optogenetic stimulation.

We injected either the control (TTT) or *Vgat* targeted viruses described above into the VP of *Vglut2*-Cre:lsl-Cas9 mice (**Figure 2A**). After 6 weeks, we performed slice electrophysiology recordings from neurons in the LHb while stimulating the VP→LHb projection terminals with blue light (1ms, 1mW/mm^2^, **Figure 2B**). We recorded oEPSCs at -55 mV and oIPSCs at +10 mV in the same cell. As expected, oEPSC amplitudes were not significantly different between edited and control groups (**Figure 2C,D**), suggesting similar efficacy of optogenetic activation with ChRonos between groups. However, we found a significant reduction in oIPSC:oEPSC ratios in edited animals compared to controls (**Figure 2C,E**), and evoked outward currents were blocked with picrotoxin (**Figure 2C**). This finding provides additional evidence for efficient *Vgat* editing in a different neuronal population with a robust internal control.

**Figure 2.**
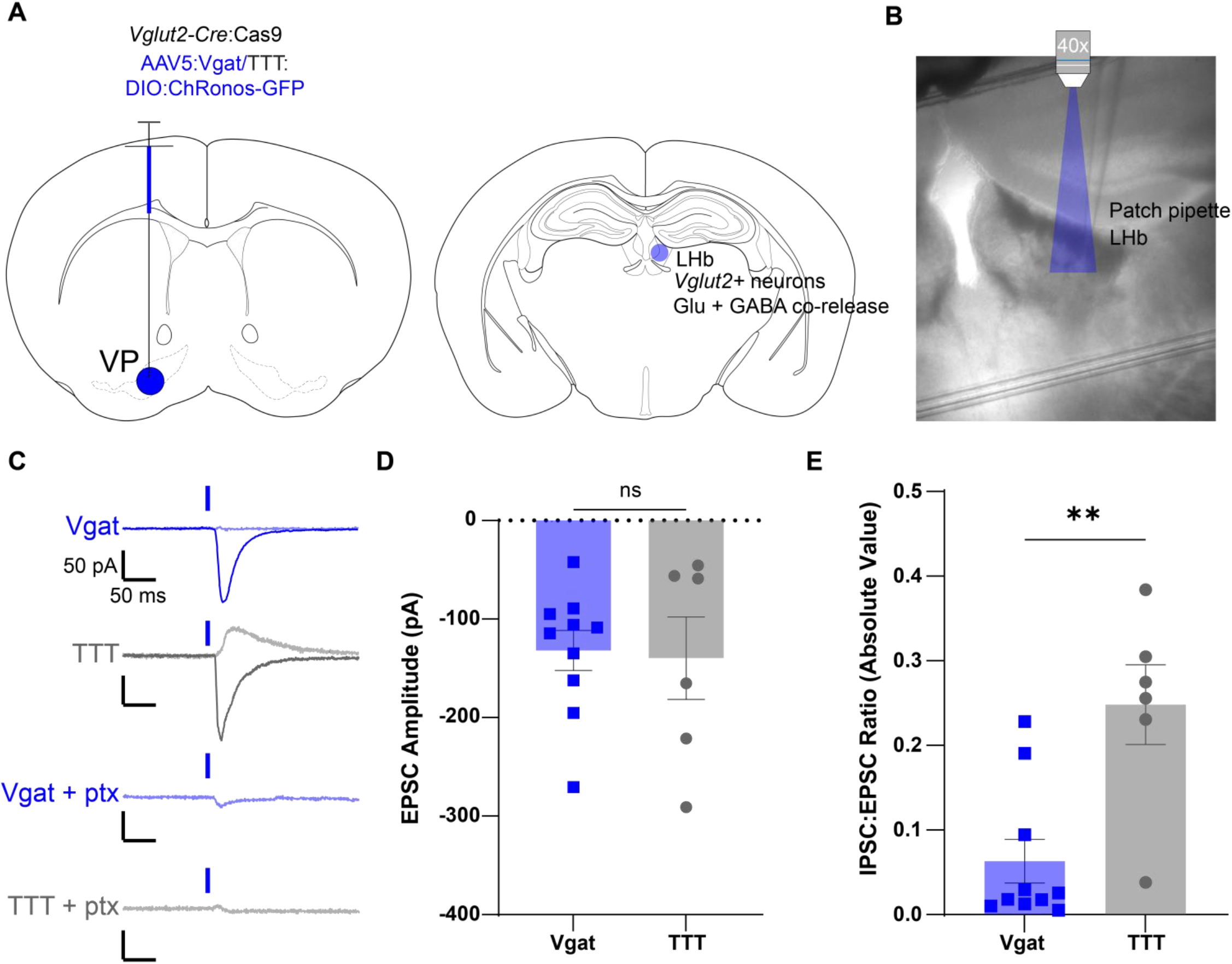
ChRonos-GFP coupled with *Vgat* editing selectively reduces inhibitory synaptic currents from GABA/Glutamate-co-releasing neurons. (**A**) Diagram of virus injection into the VP of 6-8 week *Vglut2-*Cre:Cas9 mice (left) and termination of *Vglut2*+ GABA/glutamate co-releasing neurons from the VP in the LHb (right). (**B**) Representative 4x image of slice physiology recording in the LHb. Area of blue light stimulation of ChRonos shown in blue. (**C**) Representative EPSC (solid) and IPSC (shaded) traces for Vgat-edited (1^st^ row, blue) and TTT control (2^nd^ row, grey) neurons, and representative traces for Vgat-edited (3^rd^ row, blue) and TTT control (4^th^ row, grey) oIPSC blocked with picrotoxin (ptx). Traces are aligned to light pulse onset (blue rectangle). The representative Vgat-edited oEPSC and oIPSC traces are an average of 49 consecutive sweeps each; the representative TTT control oEPSC and oIPSC traces are an average of 27 and 12 consecutive sweeps, respectively. The representative Vgat-edited oIPSC plus ptx is an average of 35 consecutive sweeps, and the representative control oIPSC plus ptx is an average of 15 consecutive sweeps. Horizontal scale bars=50 ms; vertical scale bars=50 pA. (**D**) Quantification of oEPSC amplitude for Vgat (blue) and TTT (grey) neurons. Each point represents the average evoked excitatory current amplitude from 10 ms before through 10 ms after the peak amplitude for 10-50 consecutive sweeps. The average Vgat oEPSC amplitude was -131.8±20.26 pA (n=10 neurons from 4 animals); the average TTT oEPSC amplitude was -139.6±41.84 pA (n=6 neurons from 3 animals) (p=0.87; ns). (**E**) Quantification of the absolute value of the oIPSC:oEPSC ratio for Vgat-edited (left, blue) and TTT control (right, grey) neurons. oIPSC measurements were obtained by averaging the current values 10 ms before through 10 ms after the peak current amplitude. Each point represents the absolute value of the oIPSC 20 ms average divided by the oEPSC 20 ms average. The mean Vgat oIPSC:oEPSC ratio was 0.06±0.026 (n=10 neurons from 4 animals); the mean TTT oIPSC:oEPSC ratio was 0.25±0.047 (n=6 neurons from 3 animals) (p=0.0088**). All comparisons were done using an unpaired, 2-tailed t-test with Welch’s correction for unequal variances. All data are reported as mean±SEM.

### *Grin1* editing in dopaminergic VTA neurons reduces synaptic NMDAR currents

Next, we tested whether we could use our approach to selectively record the activity of edited neurons using a fluorescent Ca^2+^ sensor. For these experiments, we targeted NMDA receptors (NMDARs) in dopaminergic neurons in the ventral tegmental area (VTA) due to robust NMDAR activation in response to rewarding stimuli (Harnett et al., 2009; Overton & Clark, 1997; Paladini & Roeper, 2014; Stuber et al., 2008; Zweifel et al., 2009). We created a viral vector containing Cre-dependent GCaMP8f (Y. Zhang et al., 2023) using the same pX552 backbone as above (**Figure 3A**). We then cloned in gRNA against the *Grin1* subunit of the NMDA receptor or a TTT control (**Figure 3B**) and packaged these constructs into AAV9 serotyped viruses. To selectively express Cas9 in dopaminergic (DA) neurons, we crossed mice with Cre knocked in to the tyrosine hydroxylase (*Th*) gene (Lindeberg et al., 2004) to a Cre-dependent Cas9 line (Chiou et al., 2015). We then injected 6-8 week old adult TH-Cre:lsl-Cas9 mice from the F1 generation with control or active virus in the ventral tegmental area (VTA) to validate NMDAR knock-down using slice electrophysiology (**Figure 3C**).

**Figure 3.**
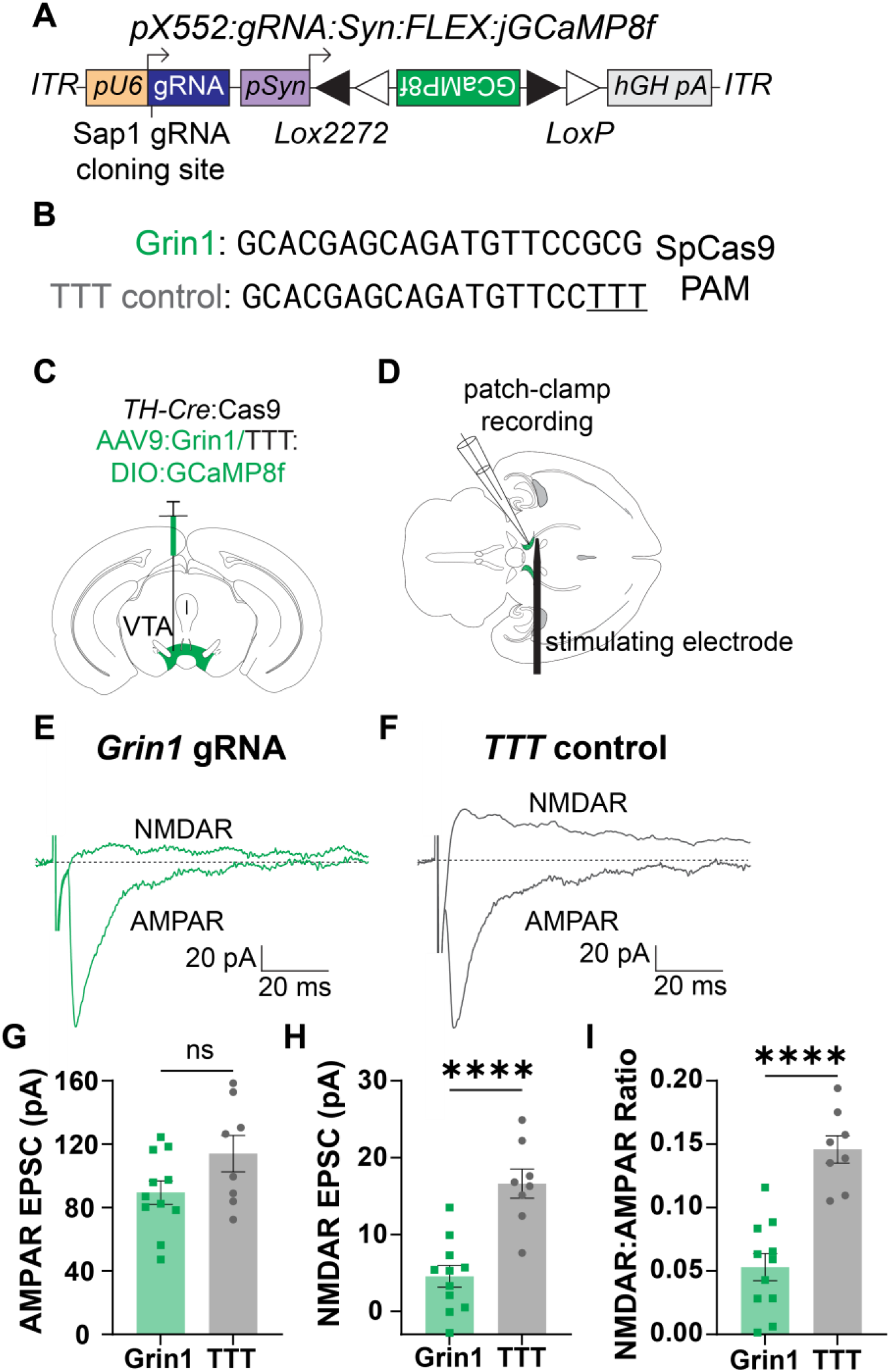
*Grin1* knockdown selectively reduces NMDAR current in VTA DA neurons. (**A**) Schematic of pX552 vector with Cre-dependent GCaMP8f transgene and Grin1 gRNA. (**B**) Sequences of gRNA targeting *Grin1* (top) and modified TTT control (bottom). (**C**) Diagram of virus injection into the VTA of 6-8 week *TH-*Cre:Cas9 mice. (**D**) Diagram of patch-clamp recording setup with recording electrode and stimulating electrode in the VTA. (**E**) Representative traces from Grin1-edited neurons of evoked NMDAR current (top) and AMPAR current (bottom). Traces are aligned to stimulating electrode pulse. Each trace represents a single trial. Horizontal scale bar=20 ms; vertical scale bar=20 pA. (**F**) Representative traces from TTT control neurons of evoked NMDAR current (top) and AMPAR current (bottom). Traces are aligned to stimulating electrode pulse. Each trace represents a single trial. Horizontal scale bar=20 ms; vertical scale bar=20 pA. (**G**) Quantification of AMPAR EPSC amplitude in Grin1 (green, left) and TTT (grey, right) neurons. Each point represents the average peak AMPAR current amplitude from 10-15 consecutive sweeps for one neuron. The average Grin1 peak AMPAR current was 89.51±7.42 pA (n=11 cells from 3 animals); the average TTT peak AMPAR current was 114.1±11.47 pA (n=8 cells from 3 animals) (p=0.096; ns). (**H**) Quantification of NMDAR EPSC amplitude in Grin1 (green, left) and TTT (grey, right) neurons. Each point represents the average peak NMDAR current amplitude from 10-15 consecutive sweeps for one neuron. The average Grin1 peak NMDAR current was 4.557±1.41 pA (n=11 cells from 3 animals); the average TTT peak NMDAR current was 16.61±1.90 pA (n=8 cells from 3 animals) (p=0.0002***). (**I**) Quantification of NMDAR:AMPAR ratio in Grin1 (green, left) and TTT (grey, right) neurons. Each point represents the ratio of the average peak NMDAR current amplitude to the average peak AMPAR current amplitude from 10-15 consecutive sweeps for one neuron. The average Grin1 NMDAR:AMPAR ratio was 0.053±0.011 (n=11 cells from 3 animals); the average TTT NMDAR:AMPAR ratio was 0.146±0.011 (n=8 cells from 3 animals) (p<0.0001****). All comparisons were done using an unpaired, 2-tailed t-test with Welch’s correction for unequal variances. All data are reported as mean±SEM.

6 weeks later, we recorded from GCaMP8f+ VTA DA neurons in acute horizontal slices and measured electrically-evoked AMPAR-and NMDAR-mediated EPSCs (**Figure 3D**). Evoked AMPAR EPSCs, measured at -70 mV, were not significantly different between edited and control animals (**Figure 3E-F, G**). However, NMDAR EPSCs were nearly abolished in *Grin1* edited neurons and significantly reduced compared to controls (**Figure 3H**). We also observed a significant reduction in the ratio of NMDAR:AMPAR currents between edited neurons and controls (**Figure 3I**). These data demonstrate that we can efficiently suppress synaptic NMDAR function in VTA DA neurons by editing *Grin1* subunits.

### In-vivo calcium imaging in gene-edited VTA neurons

Having validated the efficacy of our *Grin1* knockout using slice electrophysiology, we next combined this approach with fiber photometry to record Ca^2+^ signaling in awake, behaving animals during a conditioned Pavlovian cue/reward task. To test whether *Grin1* editing could reduce stimulus-evoked GCaMP8f Ca^2+^ signal amplitude, we injected viruses containing both gRNA against either *Grin1* or a TTT control and Cre-dependent GCaMP8f, as above, bilaterally into the VTA of adult TH-Cre/lsl:Cas9 mice (**Figure 4A**, left). We also implanted a photometry fiber above the left injection site (**Figure 4A**, right). Fiber placement was validated using histology and *in situ* hybridization for *Th* and GCaMP8f (**Figure 4B**). After 6 weeks, mice were acclimated to the recording chamber. After 3 days of acclimation, we recorded GCaMP8f Ca^2+^ transients in VTA dopaminergic neurons over a period of 5 days while the mice performed a Pavlovian cue-reward task, in which a tone accompanied the delivery of a food reward (**Figure 4C**).

**Figure 4.**
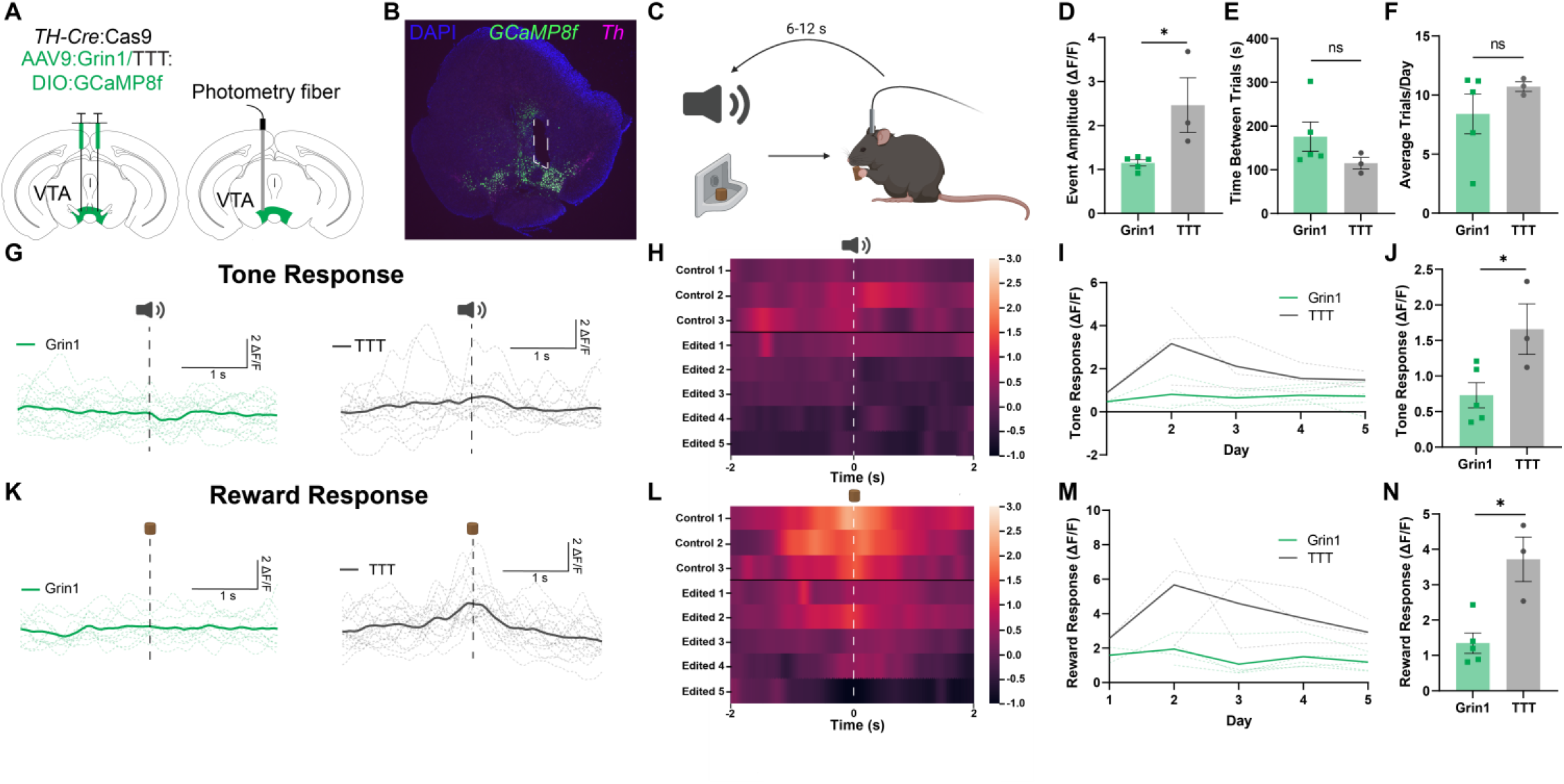
GCaMP8f Ca^2+^ currents are significantly reduced during a Pavlovian reward task after *Grin1* knockdown in VTA DA neurons. (**A**) Diagram of bilateral virus injection (left) and unilateral fiber optic insertion (right) into the VTA of 6-8 week *TH-*Cre:Cas9 mice. (**B**) Representative fluorescence ISH image of GCaMP8f virus expression in the VTA (green) overlapping with *TH*+ neurons (purple). Implant site indicated by white dashed lines. (**C**) Schematic of Pavlovian cue-reward conditioning paradigm. (**D**) Quantification of all GCaMP8f transient amplitudes in Grin1 (green, left) and TTT (grey, right) conditions. Each point represents the average event amplitude, measured as change in fluorescence over baseline (ΔF/F) across all recording sessions for one animal. The average Grin1 event amplitude was 1.15±0.07 (n=3 animals); the average TTT event amplitude was 2.46±0.62 (n=5 animals) (p=0.029*). (**E**) Quantification of time between trials for Grin1 (green, left) and TTT (grey, right) conditions. Each point represents the average time between trials across all recording sessions for one animal. The average Grin1 time between trials was 175.8±33.56 s (n=3 animals); the average TTT time between trials was 115.4±13.3 s (n=5 animals) (p=0.19; ns). (**F**) Quantification of average number of trials per day for Grin1 (green, left) and TTT (grey, right) conditions. Each point represents the average number of trials per day across all recording sessions for one animal. The average Grin1 trials/day was 8.4±1.69 trials (n=3 animals); the average TTT trials/day was 10.72±0.40 trials (n=5 animals) (p=0.25; ns). (**G**) Representative GCaMP8f response to tone cue in Grin1 (green, left) and TTT (grey, right) conditions. Dashed lines represent individual trials within a single session; solid line represents the average of all trials for a single session. Dashed black line=tone cue onset. (**H**) Heat map displaying average GCaMP8f tone response across all trial days for all animals. Dashed white line=tone cue onset. (**I**) Average peak tone cue response on each recording day for Grin1 (green) and TTT (grey) conditions. Dashed lines=daily tone response averages for individual subjects; solid lines=tone response average for Grin1 (green) and TTT (grey). (**J**) Quantification of average peak tone response for Grin1 (green, left) and TTT (grey, right). Each point represents the average peak GCaMP8f tone response across all recording sessions for one animal. The average Grin1 peak GCaMP8f tone response was 0.73±.018 (n=3 animals); the average TTT peak GCaMP8f tone response was 1.66±0.36 (n=5 animals) (p=0.038*). (**K**) Representative GCaMP8f response to reward in Grin1 (green, left) and TTT (grey, right) conditions. Dashed lines represent individual trials within a single session; solid line represents the average of all trials for a single session. Dashed black line=reward retrieval. (**L**) Heat map displaying average GCaMP8f reward response across all trial days for all animals. Dashed white line=reward retrieval. (**M**) Average peak reward response on each recording day for Grin1 (green) and TTT (grey) conditions. Dashed lines=daily reward response averages for individual subjects; solid lines=reward response average for Grin1 (green) and TTT (grey). (**N**) Quantification of average peak reward response for Grin1 (green, left) and TTT (grey, right). Each point represents the average peak GCaMP8f reward response across all recording sessions for one animal. The average Grin1 peak GCaMP8f reward response was 1.35±0.29 (n=3 animals); the average TTT peak GCaMP8f reward response was 3.72±0.63 (n=5 animals) (p=0.0085**). All comparisons were performed using an unpaired, 2-tailed, nested t-test, where each sub-column represents one animal, and each data point within the sub-column represents the average of all trials for that animal on each recording day. All data are reported as mean±SEM.

We found that, independent of task stimuli, GCaMP8f Ca^2+^ transients in edited animals were nearly absent and significantly reduced compared with control (**Figure 4D**). Upon analyzing task performance, we found that edited and control animals did not differ significantly in their time between trials (**Figure 4E**), nor in number of trials completed (**Figure 4F**). This corroborates previous reports that NMDARs in dopaminergic VTA neurons do not play a significant role in cue-reward association learning (Molina & Alvarez-Sabín, 2009; Parker et al., 2010).

Despite their similar task performance, Ca^2+^ transients in response to both cue and reward were significantly reduced in edited animals compared with controls. Edited animals displayed minimal response to the tone cue (**Figure 4G**), a trend which held across subjects (**Figure 4H**) and trial days (**Figure 4I**). Control animals, meanwhile, exhibited a modest Ca^2+^ fluorescence increase in response to the tone, and the overall average peak GCaMP8f signal in response to the tone was significantly reduced in edited animals compared with control (**Figure 4J**). The GCaMP8f transients in response to food reward were similarly reduced in edited animals (**Figure 4K**) which held across subjects (**Figure 4L**) and trial days (**Figure 4M**). Control animals, by contrast, displayed a robust food reward response. The overall average peak Ca^2+^ response to food reward was significantly reduced in edited animals compared with control (**Figure 4N**). These data show that our CRISPR/Cas9 system for gene knockout can be used in combination with fiber photometry to selectively monitor activity in genetically edited neurons during behavioral tasks.

### Anatomical tracing and CRISPR/Cas9 editing in the peripheral nervous system using systemic viral vectors

So far, we have demonstrated the efficacy of our single-vector CRISPR/Cas9 system for gene knockout in the central nervous system of adult mice. To broaden the applications of this tool, we next tested it in the peripheral nervous system of neonatal mice. Conditional knockout of *Dicer* in parvalbumin (*PV*) positive peripheral proprioceptor neurons results in axonal retraction from their downstream targets in the ventral horn of the spinal cord (Imai et al., 2016). To test whether these viral CRISPR/Cas9 approaches could be used to study genetic regulation of axonal morphology in the PNS, we first inserted a Cre-dependent mCherry transgene into the pX552 gRNA cloning vectors (**Figure 5A**). We then developed gRNAs targeted against *Dicer* or a control gRNA with mismatch sequences adjacent to the SpCas9 PAM (**Figure 5B**), cloned these into the pX552 mCherry vector, and packaged it into PNS-selective PHP.S serotyped AAVs (Chan et al., 2017). We injected active or control virus retro-orbitally in p0-1 *PV-*Cre/lsl-Cas9 mice to target PV+ proprioceptive neurons (**Figure 5C**). After 14 days, we analyzed PV+ neuronal cell bodies in the DRG using ISH to determine the infection efficiency of the virus. Both edited and control virus showed a high infection efficiency with no off-target expression (**Figure 5D, E**). Analysis of mCherry+ spinal cord projections revealed a significant reduction in both fluorescence density and area innervated by mCherry+ spinal cord projections of PV+ neurons in edited compared with control mice in the dorsal, intermediate, and ventral aspects of the spinal cord (**Figure 5F, G-I**). These findings recapitulate the phenotype seen with conditional Dicer knockout in PV+ peripheral proprioceptive neurons, with reduction in fiber density in both the ventral and dorsal horns of the spinal cord (Imai et al., 2016). These data demonstrate that our single-vector CRISPR/Cas9 system can also be used at early postnatal stages of neural circuit development in the peripheral nervous system.

**Figure 5.**
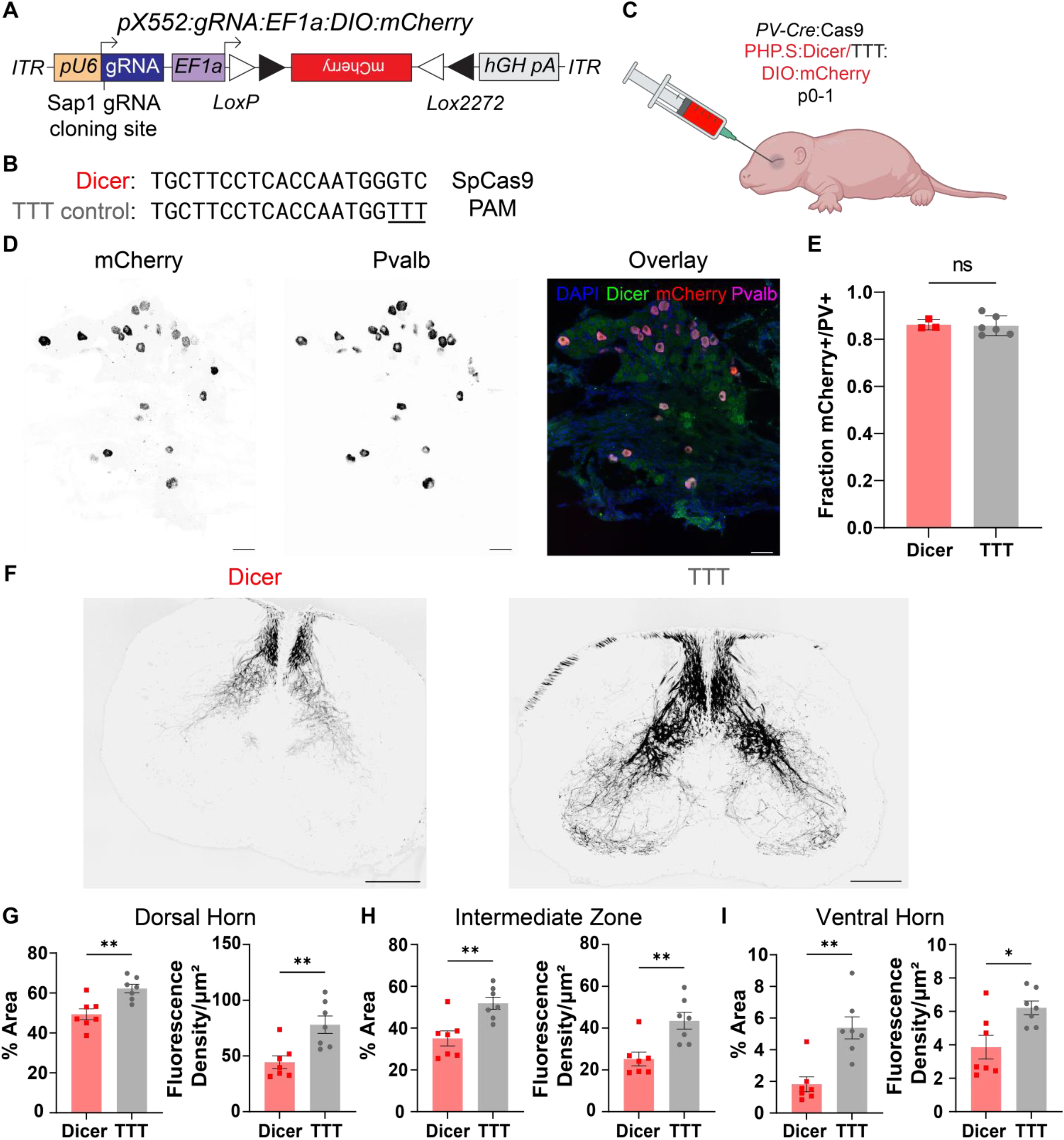
Cre-dependent mCherry expression and *Dicer* editing in peripheral PV+ neurons. (**A**) Schematic of pX552 vector with Cre-dependent mCherry transgene and Dicer gRNA. (**B**) Sequences of gRNA targeting *Dicer* (top) and modified TTT control (bottom). (**C**) Diagram of retro-orbital systemic virus injection into p0-1 *PV-*Cre:Cas9 mice. (**D**) 20X images of lumbar DRG from a p14 *PV-*Cre:Cas9 mouse labeled using ISH for *mCherry* (left) and *PV* (middle). Overlay (right) shows a high percentage of mCherry/Dicer overlap. Scale bars=50µm. (**E**) Quantification of *mCherry+*/*PV+* DRG neurons for Dicer (red, left) and TTT (grey, right). The average Dicer fraction of *PV+* cell bodies expressing *mCherry* was 0.86±0.01 (n=3); the average TTT fraction was 0.86±0.02 (n=6) (p=0.85; ns). (**F**) 4X representative images of mCherry-expressing proprioceptive neuron processes in the spinal cord for Dicer (left) and TTT control (right) groups. Scale bars=200µm. (**G**) Quantification of % Area (left) and Fluorescence density (right) of mCherry+ processes in the spinal cord dorsal horn in Dicer vs. TTT groups. The average dorsal horn % fiber area for the Dicer group was 49.34±2.76% (n=7); the average for the TTT control group was 62.26±2.13% (n=7) (p=0.003**). The average dorsal horn fluorescence density for the Dicer group was 44.47±5.66 AU/µm^2^ (n=7); the average for the TTT group was 78.08±7.80 AU/µm^2^ (n=7) (p=0.005**). (**H**) Quantification of % Area (left) and Fluorescence density (right) of mCherry+ processes in the spinal cord intermediate zone (IZ) in Dicer vs. TTT groups. The average IZ % fiber area for the Dicer group was 35.11±3.60% (n=7); the average for the TTT control group was 51.93±2.90% (n=7) (p=0.004**). The average IZ fluorescence density for the Dicer group was 25.24±3.24 AU/µm^2^ (n=7); the average for the TTT group was 43.50±3.97 AU/µm^2^ (n=7) (p=0.004**). (**I**) Quantification of % Area (left) and Fluorescence density (right) of mCherry+ processes in the spinal cord ventral horn in Dicer vs. TTT groups. The average ventral horn % fiber area for the Dicer group was 1.82±0.47% (n=7); the average for the TTT control group was 5.39±0.70% (n=7) (p=0.002**). The average ventral horn fluorescence density for the Dicer group was 3.86±0.71 AU/µm^2^ (n=7); the average for the TTT group was 6.22±0.40 AU/µm^2^ (n=7) (p=0.017*).

## Discussion

These findings demonstrate the efficacy and broad utility of our single-vector approach for CRISPR/Cas9 gene editing and expression of genetically encoded tools. This method allows for efficient gene editing in both the central and peripheral nervous system and flexible expression of a variety of genetically encoded tools to permit in-depth interrogation of target genes in specific neural circuits and brain regions. When used with optogenetic or chemogenetic techniques, this method can be a powerful tool to study mechanisms of neurotransmitter release, learning, or synaptic plasticity. Combined with *in vivo* genetically encoded biosensor imaging, it can be used to understand how specific genes influence neural circuit dynamics. And when expressed with fluorescent proteins, this tool can facilitate studies on the genetic drivers of neuronal morphology, connectivity, and survival. However, our approach has some limitations which should be taken into consideration.

Any leaky Cre expression during development will result in Cas9 positive cells that may not faithfully represent the intended cell populations in adult mice (Campsall et al., 2002; Dietrich et al., 2000; Hébert & McConnell, 2000). This should be tested with any newly developed Cre lines prior to using this approach. If this is this case, Cre-dependent Cas9 and gRNA vectors developed by the Zweifel lab (Hunker et al., 2020) would be a more effective strategy. Alternatively, future iterations of this vector design could incorporate Cre-dependent U6-gRNA cassettes to circumvent the issue of leaky Cas9 expression and ensure editing only in the cells that express Cre at the desired developmental time point. Additionally, our approach is largely limited to mice where transgenic lines are more prevalent, though Cre-dependent Cas9 transgenic rats have recently been developed (Bäck et al., 2019).

CRISPR/Cas9 gene editing is also limited by the number and arrangement of transcript variants for the target gene. This approach works best when targeting a gene with one or a few transcript variants which all share at least one common exon, preferably early in its coding sequence (Hunker et al., 2020; Joung et al., 2017; Shalem et al., 2014). It is more challenging to effectively target genes for which there are many transcript variants with variable exon structures. Future improvements on this approach could involve introducing multiple gRNAs in the same vector, allowing for better targeting of all variants. By using Cre-dependent Cas9 mouse lines, rather than packaging Cas9 in the viral vector, there is ample space remaining in the virus for additional gRNAs to facilitate improved targeting of a single gene or targeting of multiple genes simultaneously.

Additionally, CRISPR/Cas9 editing may not completely knock out the target gene, rather, it functions more as a “knock-down.” In our slice electrophysiology recordings in the NAc, for example, we noted residual evoked IPSCs, even in areas of strong virus expression. This could be a result of the randomness in relying on NHEJ repair to introduce frameshift mutations, and some neurons may have retained at least one functional copy of the target gene (Canver et al., 2014; Hunker et al., 2020). This could also be due to retention of existing VGAT that was produced prior to gene editing, or non-specific transport of GABA into vesicles via other solute transporters (Tritsch et al., 2012). These limitations highlight the importance of validating knock-down efficacy for each gene and model system.

In *PV*+ spinal cord projections of *Dicer*-edited mice, we were able to recapitulate the phenotype of reduced innervation seen with conditional Dicer knockout (Imai et al., 2016). However, we noted a small population of persistent *PV+* projections in the ventral horn. This could again be due to incomplete editing of *Dicer* in all infected *PV*+ neurons. Alternatively, it is possible that some percentage of *PV*+ proprioceptive neurons have already become synaptically anchored in the ventral spinal cord by the time that we injected virus at p0-1, and thus were less susceptible to retraction due to *Dicer* editing. This possibility highlights one benefit of our approach over traditional conditional knockout: the ability to precisely time editing to probe the role of genes of interest during development. This feature could be used to edit key genes involved in neuronal migration or synapse formation at multiple time points to better understand the precise developmental stages at which these genes function. Future studies could also use our gene editing approach in combination with *in utero* electroporation or viral transduction to achieve targeted gene editing at even earlier developmental time points.

This CRISPR/Cas9 approach could be used to more efficiently screen for the effects of editing gene candidates identified through clinical GWAS studies or transcriptomic studies from defined cell types in model organisms. Though not included here, these vectors could also be modified to express other Cre-dependent opto- or chemogenetic approaches to enable further mechanistic understanding of various genes throughout the nervous system. Our approach is also not limited to targeting neurons; in combination with the appropriate viral vectors and mouse lines, this tool could be used to manipulate genes in astrocytes or cells in other organs. Finally, this tool could be used to knock-down native gene expression and re-introduce a mutant gene variant in defined cell types, increasing its translational potential to study the impacts of mutations identified in clinical settings.

## Conclusion

In this study, we present a flexible, single-vector approach to CRISPR/Cas9 editing and expression of genetically encoded tools. We demonstrate its utility in various cell types and brain regions throughout the central nervous system and describe the first systemic CRISPR/Cas9 gene editing with co-expressed reporters in the PNS. By combining our single-vector approach with genetic tools commonly used in neuroscience—channelrhodopsins, GCaMP, and fluorescent proteins—we demonstrate its potential for flexible and precise interrogation of gene function in specific cell types and circuits throughout the nervous system.

## Acknowledgements

This work was supported by NIH R01s NS130046 (BAC), DA049924 (MCC), DA058755 (MCC), and internal funds from the McDonnell Center for Systems Neuroscience (BAC). We would like to thank all members of the lab for helpful discussions and feedback. We thank Alexxai Kravitz for his feedback on our photometry experiment design. We thank Alex Legaria for help with photometry analysis. We would also like to thank Dr. Mingjie Li at the Hope Center Viral Vectors Core at Washington University for assistance in establishing viral vector purifications. Finally, we would like to thank Vera Thornton for consulting on statistical analyses. AAV PHP.S viruses were purchased from the UNC Neuroscience Center/BRAIN Initiative NeuroTools Core (U24 NS124025 to Kimberly Ritola).

## Author Contributions

JCM, MCC and BAC designed the study. JCM and BAC wrote the manuscript with input from all authors. JCM designed and cloned gRNAs, produced viruses, conducted and analyzed all photometry and behavioral experiments, and analyzed all histology data. JCM, INB, and BAC performed virus injections and processed tissues for histology. VK generated all viral vectors.

JRT, MCC and BAC performed slice electrophysiology experiments. MH maintained the transgenic mouse lines and oversaw colony maintenance. MCC and BAC supervised the project and acquired funding.

## Methods

### Lead contact

Further information and requestions for resources and reagents should be directed to the Lead Contact Byran Copits.

### Material availability

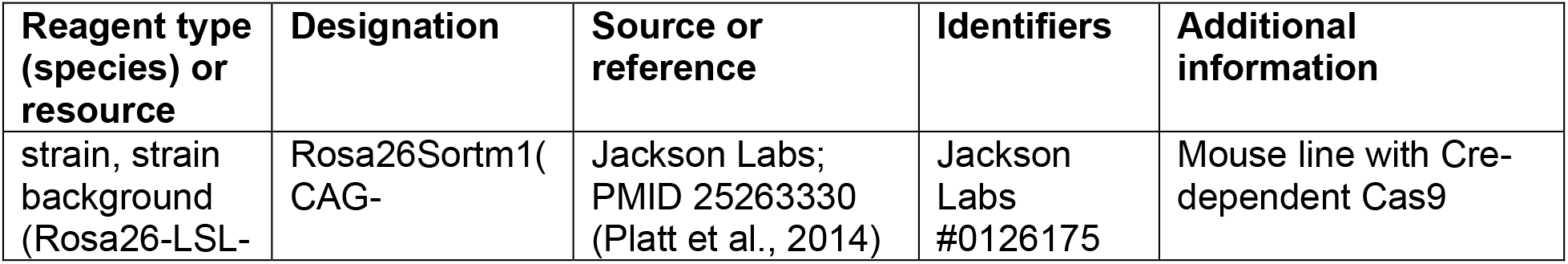
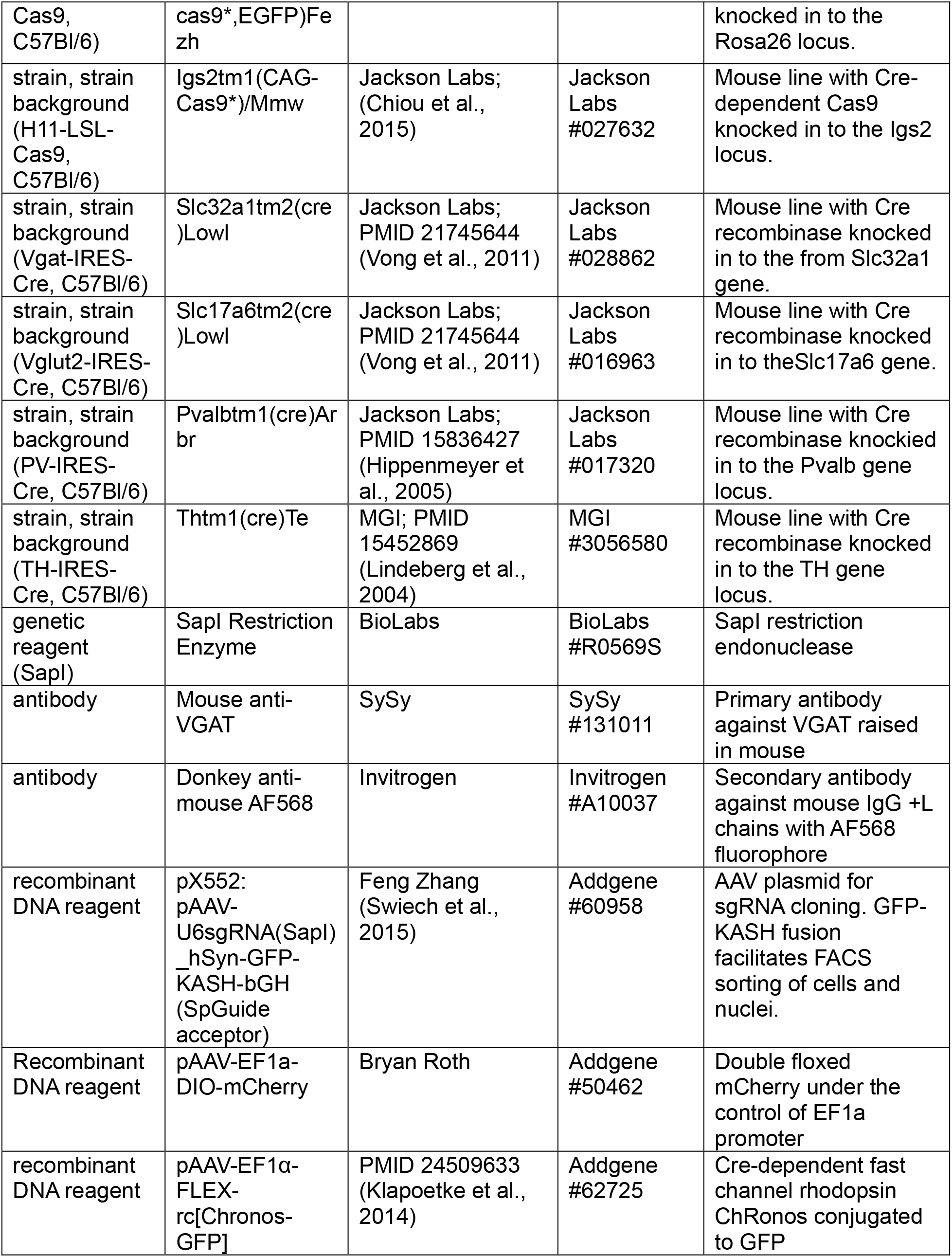
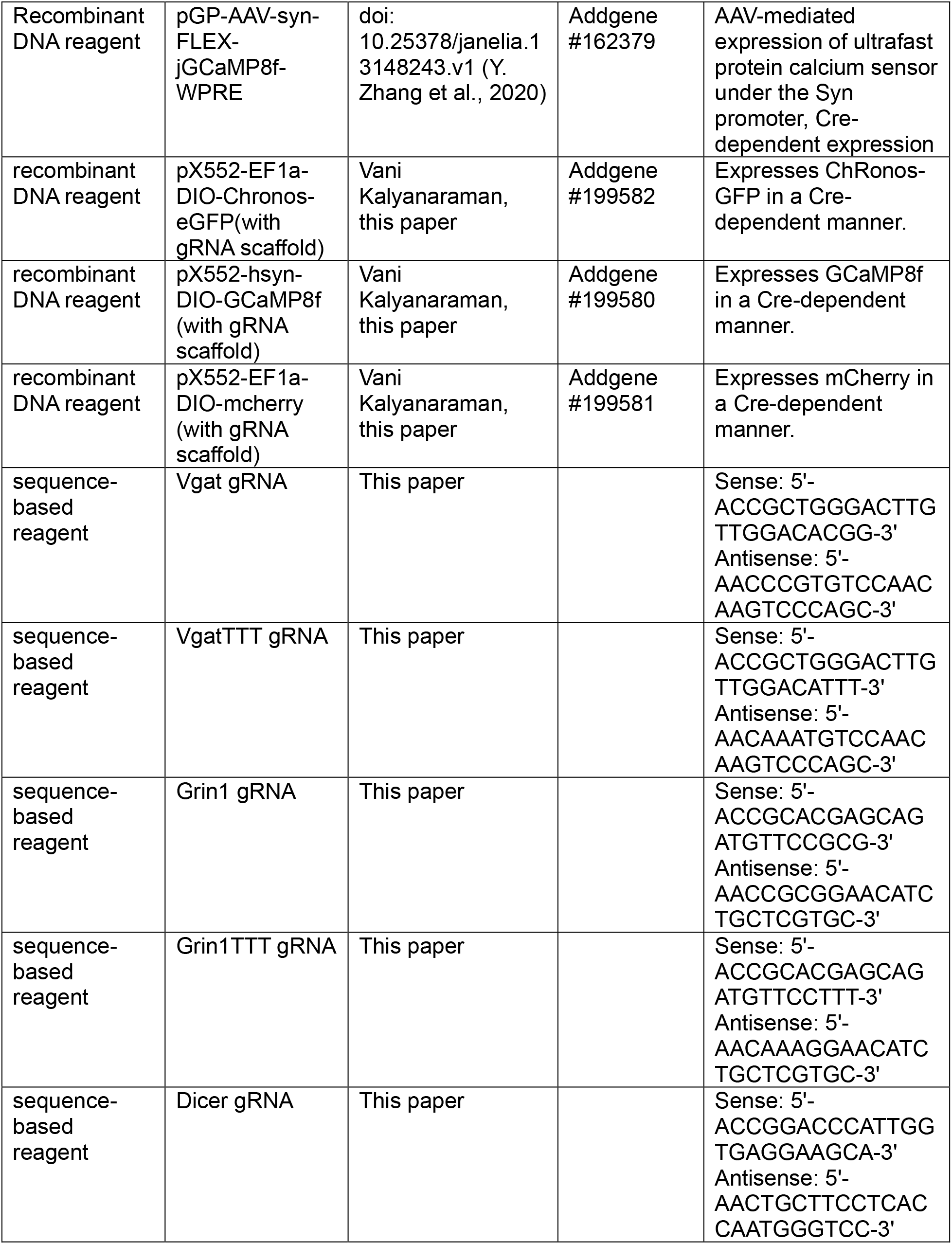
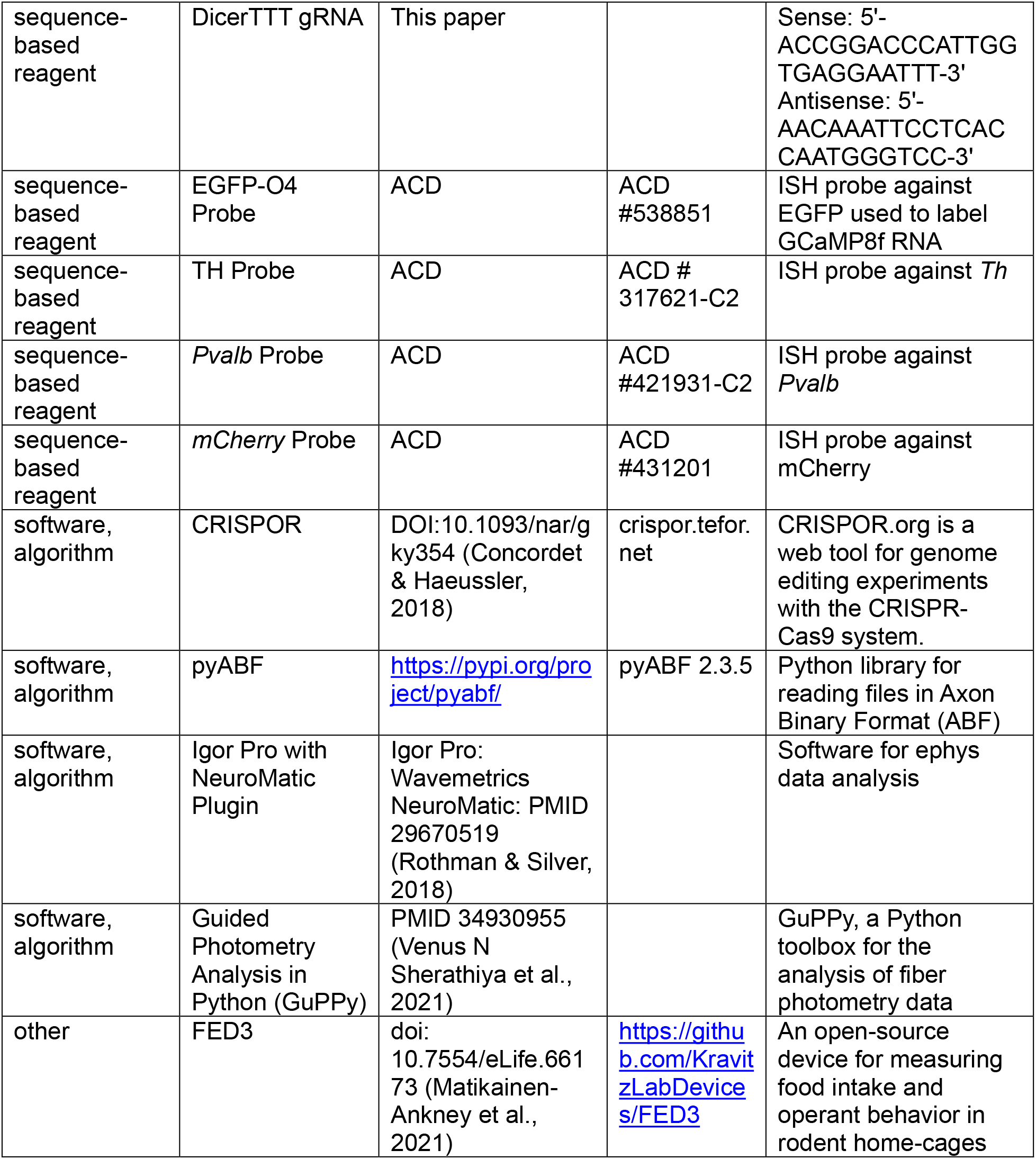

All DNA and viral constructs generated here are deposited on Addgene and are also available upon request from the Lead Contact.

### Data and code availability

All data and analysis codes are available from the Lead Contact.

### Viral vector generation

#### Cloning Cre-dependent transgene into pX552 empty vector

For all vectors, the pX552 vector (Addgene #60958) (Swiech et al., 2015) with gRNA scaffold was used as a backbone.

DIO-mCherry in pX552: Cut pcDH-DIO-mCherry with EF1a promoter with EcoRV and EcoR1. Cut pX552 with PspOM1, fill and cut with EcoR1.

DIO-Chronos-GFP: Cut pX552-EF1a-DIO-mcherry with SalI, fill and cut with HindIII. Cut pAAV-EF1a-FLEX-rChronos-GFP with EcoR1, fill and cut with HindIII.

To make DIO GCamp8f, we cut pX552-DIO-Synaptophysin-GFP with SpeI and EcoRV and made a G block for GCamp8f without SapI (since it has an internal SapI) with the same two enzymes. Ligate and transform.

#### Selecting gRNA candidates

To select candidate gRNA sequences (Hunker et al., 2020) for each target gene, we first used the National Center for Biotechnology Information (NCBI) Gene database to assess the exon structure and compare the splice variants of the target gene. The earliest exon shared among all splice variants was used as the target region for the gRNA search. We then obtained the coding sequence for the target exon from Ensembl.org. We used the CRISPOR web tool to identify suitable gRNA sequences within the target exon adjacent to *Streptococcus pyogenes* Cas9 Protospacer Adjacent Motifs (spPAMs) (Concordet & Haeussler, 2018). We pasted the exon coding sequence into the text box below ‘Step 1’ on the CRISPOR home page, selected the ‘Mus musculus – Mouse (reference) – UCSC Dec. 2011 (mm10=C57BL/6J)’ under ‘Step 2’, and selected ‘20bp-NGG – SpCas9, SpCas9-HF1, eSpCas9 1.1’ under ‘Step 3’. Two candidate sequences for each gene were selected that maximized the Doench ’16 Predicted Efficiency (Doench et al., 2016) and MIT and CFD Specificity Scores while minimizing off-target matches within the mouse genome.

Sense and antisense candidate gRNA oligos were ordered from Genewiz/Azenta with the following modifications: (1) if the original gRNA candidate did not have a ‘G’ residue at the 5’ end, a ‘G’ was added to the 5’ end of the sense oligo, with a complimentary ‘C’ added to the 3’ end of the antisense oligo. (2) To facilitate integration into Sap1 cut site sticky ends, the sequence ‘ACC’ was added to the 5’ end of the sense oligo, and the sequence ‘AAC’ was added to the 5’ end of the antisense oligo.

#### Cloning gRNA into pX552 gRNA site

2 ug of pX552 vector with SapI gRNA insertion site and Cre-dependent transgene was digested with 1:100 SapI enzyme (BioLabs #R0569S) in 1:10 CutSmart Buffer (BioLabs #B6004S) at 37°C for 1 hour. Following the first incubation, 2 µL FastAP (Thermo Scientific #EF0651) was added to the solution, which was incubated at 37°C for 1 hour, then 65°C for 20 minutes.

During the second digest, a 0.8% low-melt agarose gel with 50 µL combs was prepared. Sybr Safe dye (Invitrogen #S33102) was added to 5X DNA loading buffer at a ratio of 1 µL dye:50 µL loading buffer. Following incubation, 10.5 µL of 5X loading buffer/dye mixture was added to the 42 µL digest, which was run alongside a 1kb ladder on 0.8% agarose gel at 80V for 30 minutes. As a quality control, 2 µg of undigested vector was run alongside the digested product. After 30 minutes, gels were subjected to blue light and assessed for a ∼5 kb single band of linear DNA. This band was cut from the gel and digested using the Takara Gel Purification kit (Takara #740609.50).

Meanwhile, 10 µL phosphorylation mix (1µL 100mM sense oligo, 1µL 100mM anti-sense oligo, 0.5µL 25mM ATP (BioLabs #P0756S), 1µL 10X PNK T4 Buffer (BioLabs #B0201S), 1µL T4 PNK (BioLabs #M0201L), 5.5µL dH_2_O) was prepared. gRNA oligos were phosphorylated according to the following protocol: (1) 37°C for 30 minutes; (2) 95°C for 5 minutes; (3) Cool to 4°C at 0.1°C/sec. Phosphorylated oligos were diluted 1 µL oligo:49 µL dH_2_O. Following phosphorylation, 10µL ligation mix was prepared (20ng digested pX552 vector, 0.8µL 1:50 diluted oligo mix, 5µL 2X T7 Buffer (BioLabs #B0318S), 0.5µL T7 DNA Ligase (BioLabs #M0318S) in dH_2_O. Vectors were ligated at room temperature for 30 minutes.

Following ligation, 5 µL ligated vector was added to 25 µL Stbl3 cells (Invitrogen #C737303) and mixed by gently tapping. The mixture was incubated on ice for 30 minutes, then heat-shocked in a 42°C water bath for 45 seconds. After heat shock, cells were placed on ice for 2 minutes before adding 250 µL SOC medium (Biolabs #B9020S). The transformed cells in medium were incubated in a shaker for 1 hour at 37°C and 225 RPM. 50 µL transformed cells in SOC medium were plated on agar plates with 1:1000 carbenicillin (Sigma #C1389) and incubated overnight at 37°C. Following overnight incubation, 3 colonies of each construct were selected and incubated in 4 mL LB medium (Sigma #L3022) with 1:1000 carbenicillin overnight in a shaker at 37°C and 250 RPM. DNA from expanded colonies was purified using QIAprep Spin Miniprep Kit (Qiagen #27106) and concentration was determined using an Invitrogen Qubit fluorometer (Invitrogen #Q32857) and dsDNA Quantification Assay Kit (Invitrogen #Q32850). Samples were sent to Genewiz/Azenta to be sequenced from the U6 promoter. Sample sequences were assessed for integration of the correct gRNA sequence and for any replication errors in the surrounding plasmid sequence. One sample of each construct was expanded overnight in 200 mL LB with 1:1000 carbenicillin at 37°C and 250 RPM. DNA from expanded colonies was purified using the Promega Midiprep kit (Promega #A7640) and concentration was determined using a Qubit fluorometer. A sample of the final purified plasmid with gRNA insert was sent to Genewiz/Azenta for AAV-ITR sequencing of the entire packaging region.

### AAV Production

#### HEK Cell Transfection

All AAV production protocols were performed as previously described (Challis et al., 2019). One day prior to transfection, HEK293T cells were split into three 15 cm tissue culture-treated petri dishes (Corning #353025) and incubated in a 37°C tissue culture incubator with 5% CO_2_ overnight. PEI (Polysciences #239661) for transfection was prepared by dissolving 100 mg PEI in 310 mL H_2_O and titrating the solution to pH 3. PEI solution was then incubated at 37°C for 4 hours, shaking vigorously every half-hour. Finally, the solution was incubated overnight at 37°C, filter-purified the next day and stored at -20°C.

For transfection, we first prepared master mix of PEI in 1X DPBS (483µL PEI per 1000µL total volume). 1000 µL of master mix is required per 15 cm plate, so we prepared 3000 µL PEI+PBS master mix. We then prepared 3000µL DNA master mix in 1X DPBS (17.11µg pX552, 68.45µg AAV capsid, 34.23µg pHelper). 3000 µL PEI+DPBS master mix was then added to the DNA+DPBS master mix dropwise, mixed by inverting 5x, incubated at room temperature for no more than 10 minutes, and inverted 5x again. 2000 uL of the combined mixture was added to each plate of HEK cells; plates were returned to the incubator for 72 hours.

#### HEK Cell Harvesting and Stock Solution Preparation

After 72 hours, transfected HEK cells were harvested by pouring off the growth media and adding 5 mL DBPS per plate, then HEK cells were detached using a cell scraper. Cells suspended in DPBS from each of the 3 plates were transferred to a 50 mL conical tube and centrifuged at 2000 xg and room temperature for 15 minutes. Supernatant was discarded, and cell pellets were stored at -80°C overnight.

Next, we prepared solutions for AAV extraction. Lysis buffer (150 mM NaCl, 50 mM Tris Base) was prepared in ddH_2_O and titrated to pH 8.5 using 12N HCl. A stock solution of 5M NaCl in ddH_2_O and 5X PBS-MK (1mM MgCl_2_, 2.5mM KCl in 5X PBS) were also prepared. Finally, 15%, 25%, 40%, and 60% Iodixanol (IOD) stock solutions were prepared (**15% IOD:** 30mL 60% IOD, 24mL 5M NaCl, 24 mL 5X PBS-MK, 42mL ddH_2_O; **25% IOD:** 33mL 60% IOD, 16mL 5X PBS-MK, 0.2mL Phenol Red, 30.8mL ddH_2_O; **40% IOD:** 80mL 60% IOD, 24mL 5X PBS-MK, 16mL ddH_2_O; **60% IOD:** 80mL 60% IOD, 0.2mL Phenol Red).

#### AAV Extraction and Purification

The next day, frozen HEK cell pellets were thawed in a 37°C water bath. 5 mL lysis buffer was added to the thawed pellet and vortexed until the pellet was completely resuspended. The lysis buffer suspension was then subjected to three freeze-thaw cycles at -80°C and 37°C, respectively. After the final thaw, 14.46 µL of 1M MgCl2 and 5 µL Benzonase were added to the solution, which was then vortexed and incubated at 37°C for 30 minutes. Following incubation, the solution was vortexed again and centrifuged at 350 xg and 4°C for 20 minutes.

During centrifugation, iodixanol gradients were poured in 29.9mL Opti-seal ultracentrifuge tubes (Beckman Coulter #361625). Gradients were poured in the following order: (1) 7.5mL 15% IOD, (2) 5mL 25% IOD, (3) 7.5mL 40% IOD, (4) 5mL 60% IOD by pressing the tip of the serological pipette against the bottom of the ultracentrifuge tube and releasing the contents slowly, so as not to disrupt the gradient. Each subsequent solution was layered under the previous, and care was taken to avoid air bubbles, which can disrupt the gradient layers.

After centrifugation, supernatant was collected from the 50 mL conical tubes using a 2 mL serological pipette and slowly layered on top of the iodixanol gradient to avoid mixing with the 15% layer. The pellet of HEK cell debris was discarded. After all supernatant was added to the tube, all ultracentrifuge tubes were weighed and balanced with additional lysis buffer as needed. Tubes were capped and dried before being placed in a pre-chilled Ti-70 rotor, ensuring the tubes are placed in a balanced configuration. Tube spacers were placed on top of each tube, and the gradients were centrifuged at 350,000xg and 18°C for 1.5 hours.

During ultracentrifugation, we prepared Cytiva Vivaspin 20 100kDa MWCO concentrator tubes (Cytiva #28932363) to concentrate the final product. 5 mL 70% ethanol was added to each tube and centrifuged at 4000 rpm for 5 minutes. After centrifugation, we discarded the ethanol and added 10 mL ddH2O to the tube, inverted 5 times, and discarded the water. Another 10 mL ddH2O was added to the tube and centrifuged at 4000 rpm for 5 minutes, then discarded. After preparing the concentrator tubes, we prepared a stock solution of DPBS plus 1:1000 Pluronic detergent (Gibco #24040032) to prevent virus adhesion to the tube walls. We removed the plunger from a 10 mL syringe and attached a 0.22 µm filter (TPP #99722) to the end, then added 10 mL DPBS-Pluronic mixture to the syringe barrel. We allowed the solution to drip via gravity into the prepared concentrator tubes, timed such that there is 2-5 mL DPBS left in the syringe barrel by the time the iodixanol + virus is ready to be added.

After ultracentrifugation, tubes were removed from the rotor and placed in a tube rack. Tube caps were removed, and virus was extracted from the 40%/60% Iodixanol layer through the side of the tube using an 18-gauge needle attached to a 10 mL syringe. The needle was inserted through the tube bevel-up just above the 60% layer. When approaching the 40%/25% interface, the needle bevel was rotated down to avoid extracting the protein layer. In total, approximately 5-7 mL of iodixanol/virus mixture is extracted per virus preparation. Virus/iodixanol mixture was then layered below the DPBS/Pluronic solution described above. We inserted the plungers back into the 10 mL syringes and pushed the iodixanol+virus solution through the filter and into the upper chamber of the concentrator tube. Concentrators were then centrifuged at 3,000 xg at room temperature for 8 minutes. We repeated centrifugation until 2-5 mL of iodixanol/virus mixture remained in the upper chamber. At that point, flow-through in the bottom chamber was discarded, and 13 mL DPBS/Pluronic solution was added to the upper chamber and mixed via pipetting. We continued centrifugation at 3000 xg and room temperature until 200-500 µL of DPBS+virus remained in the upper chamber. Virus was removed from the upper chamber of the concentrator and stored in low protein binding tubes (Eppendorf #022431064) at -80°C.

After purification, AAV stock concentration was determined using the Takara AAV Titration kit (Takara #6233) and an Applied Biosystems 7500 Fast Real-Time PCR System.

### Model organisms

All procedures were conducted in accordance with National Institutes of Health guidelines and with approval from the Institutional Animal Care and Use Committee at Washington University School of Medicine. To generate transgenic lines with cell-specific expression of SpCas9, homozygous Rosa26-LSL-Cas9 knock-in mice (Rosa26Sor^tm1(CAG-cas9*,EGFP)Fezh^, Jackson labs #0126175) (Platt et al., 2014) were crossed to following homozygous Cre lines: Vgat-IRES-Cre (Slc32a1^tm2(cre)Lowl^, Jackson labs #028862) (Vong et al., 2011), Vglut2-IRES-Cre (Slc17a6^tm2(cre)Lowl^; Jackson labs #016963) (Vong et al., 2011). Homozygous H11-LSL-Cas9 knock-in mice (Igs2^tm1(CAG-Cas9*)/Mmw^; Jackson labs #027632) (Chiou et al., 2015) were crossed to homozygous TH-IRES-Cre (Th^tm1(cre)Te^, MGI 3056580) (Lindeberg et al., 2004), or PV-IRES-Cre (Pvalb^tm1(cre)Arbr^; Jackson labs #017320) (Hippenmeyer et al., 2005). All lines were maintained on a C57Bl/6 background and we used mice heterozygous for both Cas9 and Cre recombinase for all experiments. All animals were group housed with *ad libitum* access to food and water and maintained on a 12 hour light:dark cycle. We used both male and female mice for all experiments and did not observe any effects of sex in our analyses.

### Stereotaxic Surgery

For all intracranial virus injections, mice were initially anesthetized in an induction chamber with 5% Isoflurane (Piramal Critical Care #6679401710) in room air. Following induction, their heads were shaved and disinfected with iodine and 70% ethanol, and they were secured in the stereotax using ear bars. We injected mice with 0.05 mg/kg of 1 mg/mL Buprenorphine ER (ZooPharm) and a bolus of 1 mL 0.9% sterile saline (Pfizer #00409488850) at the beginning of the surgery. A midline incision was made on the scalp to expose the skull, and the head was balanced left to right and front to back using bregma as the reference coordinate. All injections were made using a 32 gauge 1.0 µL Hamilton Neuros Syringe Point Style 4 (Hamilton #65458-02). Virus was infused at a rate of 100 nL/min, and the syringe was left in place for 10 minutes after infusion.

#### NAc

We injected 6–8-week-old Vgat-IRES-Cre/Rosa26-LSL-Cas9 mice with 150 nL of AAV5-Vgat-DIO-ChRonos-GFP (active) or AAV5-VgatTTT-DIO-ChRonos-GFP (control) at 3 coordinates along the dorsal/ventral axis: A/P +1.7mm; M/L 0.8mm (left); D/V -4.65, -4.5, -4.4mm, for a total of 450 nL virus injected per animal. Both control and active virus were titered to 1×10^12^ vg/mL.

#### VP

We injected 6-8 week old Vglut2-IRES-Cre/Rosa26-LSL-Cas9 mice with 500 nL of the above Vgat or VgatTTT viruses at the following coordinates: A/P +0.16mm; M/L 1.5 mm (left); D/V -4.6 mm.

#### VTA

We injected 6-8 week old TH-IRES-Cre/H11-LSL-Cas9 mice with 1000 nL of either AAV9-Grin1-DIO-GCaMP8f (active) or AAV9-Grin1TTT-DIO-GCaMP8f (control) at the following coordinates: A/P -3.1mm; M/L 0.5mm; D/V -4.5mm. For animals used for electrophysiology, injections were performed unilaterally on the left side. For animals used for fiber photometry, the skull was scored before virus injection. Injections were performed bilaterally, and a fiber optic probe (Doric #B28044015) was inserted unilaterally on the left side during the same surgery at A/P -3.1mm; M/L 0.5mm; D/V -4.55mm. Both control and active virus were titered to 4×10^11^ vg/mL. For these animals, the fiber optic was secured in place first using C&B Metabond (Parkell #S380), then Jet Denture Repair Acrylic (Lang #1223).

### Retro-Orbital AAV Injection

To target peripheral proprioceptive neurons, we injected p0-1 PV-IRES-Cre/H11-LSL-Cas9 mice retro-orbitally with 1×10^12^ vg of either PHP.S-pX552-Dicer-DIO-mCherry (active) or PHP.S-pX552-DicerTTT-DIO-mCherry (control) (packaged by UNC Neuroscience Center/BRAIN Initiative NeuroTools Core). Injections were done using a BD Ultra-Fine insulin syringe (BD #328440) following previously established protocol (Yardeni et al., 2011).

### Electrophysiology

#### Acute Slice Preparation

Brain slices for electrophysiology recordings were prepared using a protective cutting and recovery method (Copits et al., 2021; Ting et al., 2014). For NAc recordings, 6 weeks after viral injections mice were deeply anesthetized with ketamine and xylazine and transcardially perfused with cold oxygenated NMDG-aCSF containing (in mM): 93 N-methyl-D-glucamine, 2.5 KCl, 1.25 NaH2PO4, 30 NaHCO3, 20 HEPES, 25 glucose, 5 ascorbic acid, 2 thiourea, 3 sodium pyruvate, 0.5 CaCl2, 5 MgCl2, pH=7.3, 300–310 mOsm. For VTA recordings, 6 weeks after viral injections mice were deeply anesthetized with ketamine and xylazine and transcardially perfused with cold oxygenated choline aCSF containing (in mM): 93 choline chloride, 2.5 KCl, 1.25 NaH2PO4, 30 NaHCO3, 20 HEPES, 25 glucose, 5 ascorbic acid, 2 thiourea, 3 sodium pyruvate, 12 N-acetyl-L-cysteine, 0.5 CaCl2, 5 MgCl2, pH=7.3 300–310 mOsm. Brains were rapidly removed, embedded in 2% low-melt agarose (Sigma, A0676) and 200 μm thick horizontal (VTA) or 300 μm thick coronal slices (NAc) were cut using a Compresstome (Precisionary Instruments, cat. # VF210–0Z). Slices were transferred to a recovery chamber containing oxygenated choline aCSF (VTA) or oxygenated NMDG-aCSF (NAc) at 32°C for 10 minutes before transfer to a holding chamber filled with oxygenated aCSF at 32°C containing (in mM): 124 NaCl, 2.5 KCl, 1.25 NaH2PO4, 24 NaHCO3, 5 HEPES, 12.5 glucose, 2 CaCl2, 1 MgCl2, pH=7.3, 300–310 mOsm. Slices were maintained in the dark at room temperature and allowed to recover >1 hour before recording.

#### NAc IPSC recordings

Slices were transferred to the chamber of an upright microscope (Slicescope, Scientifica), perfused with room temperature oxygenated aCSF described above at ∼2 ml/min, and visualized using IR-DIC microscopy. Evoked inhibitory postsynaptic currents (IPSCs) were recorded at 0 mV from GFP-negative cells of the nucleus accumbens in a field of Chronos+ terminals from adjacent neurons. Whole-cell recordings were made using 4–5 MΩ pipettes filled with Cs^+^ internal solution, containing (in mM) 110 cesium gluconate, 8 tetraethylammonium chloride, 3 QX-314 bromide, 1.1 EFTA, 0.1 CaCl_2_, 10 HEPES, 4 MgATP, 0.4 Na_2_GTP, 10 Na_2_phosphocreatine, pH to 7.28 with CsOH, 292 mOsm. IPSCs were pharmacologically isolated with 10 μM NBQX and 50 μM D-APV (both from HelloBio). Recordings were performed using pClamp11 software (Molecular Devices) controlling a Multiclamp 700B amplifier. Data were digitized at 10 kHz and filtered at 3 kHz. Cells were discarded if the series resistance was >30 MΩ or changed more than 20%. Wide-field photostimulation was delivered through a 40x objective using custom LEDs coupled to the back fluorescence port of the microscope (Copits et al., 2021). LEDs were triggered by TTL pulses from the amplifier to an LED current controller (Thorlabs, DC4104). All light intensities were calibrated using a photodiode (Thorlabs, S121C) and power meter (Thorlabs, PM100D).

#### LHb evoked IPSC and EPSC recordings

Whole-cell patch-clamp recordings were made from coronal slices (220 µm) of the LHb. Slices were prepared using a vibratome (Leica VT 2100) in ice-cold cutting solution (in mM: 0.5 CaCl2, 110 C5H14CINO, 25 C6H12O6, 25 NaHCO3, 7 MgCl2, 11.6 C6H8O6, 3.1 C3H3NaO3, 2.5 KCl and 1.25 NaH2PO4) and continuously bubbled with 95% O2 and 5% CO2. Slices were incubated at 32°C for 20 min in artificial cerebrospinal fluid (aCSF) (in mM: 119 NaCl, 2.5 KCl, 1.3 MgCl2, 2.5 CaCl2, 1.0 Na2HPO4, 26.2 NaHCO3 and 11 glucose) followed by storage at room temperature until electrophysiological recordings were performed. Slices were hemisected and superfused with aCSF at 30 ± 2°C. Neurons were recorded using borosilicate glass pipettes (4–5 MΩ resistance) pulled on a micropipette puller (Narishige PC-100) filled with cesium-based internal solution (in mM: 135 cesium methanesulfonate, 10 potassium chloride, 10 HEPES, 1 Magnesium chloride, 0.2 EGTA, 4 Mg-ATP, 0.3 GTP, 20 phosphocreatine, at pH 7.3 and 289 Osm). Neurons were held at -55 mV to assess glutamatergic synaptic transmission, and +10 mV to assess GABAergic synaptic transmission. Currents were amplified, filtered at 2 kHz and digitized at 10 kHz using a MultiClamp 700B amplifier and Digidata 1550 (Molecular Devices). Clampex version 11.4 (Molecular Devices) was used for data acquisition. Series resistance was monitored using a hyperpolarizing step of −15 mV for 5 ms every 10 s; the cell was discarded if the series resistance changed by more than 15%. Neurons were visualized using an Olympus ×560 upright microscope; field LED illumination (CoolLED) was used to visualize and stimulate channelrhodopsin-expressing terminals (473 nm, paired 4 ms light pulses, 50 ms interstimulus interval, 13.7–18.2 mW). 100 μM picrotoxin was washed on to abolish GABAergic transmission. All agents were purchased from Sigma Biosciences.

#### VTA NMDAR/AMPAR Recordings

Recordings were performed as above for the NAc. GCaMP8f+ neurons were identified using epifluorescence and AMPA and NMDAR currents were recorded at -70 and +40 mV, respectively. A bipolar stimulating electrode (FHC cat. #30202) connected to a stimulus isolator (WPI cat. #A365) was placed ∼200 μm rostral to the recorded cell to deliver electrical stimulation (0.5 ms). EPSCs were pharmacologically isolated with 10 μM bicuculline and 100 μM picrotoxin (both from HelloBio).

### Electrophysiology Analysis

#### NAc evoked IPSC analysis

Evoked IPSC recordings were exported in Axon Binary File (.abf) format. We used the pyABF package (Harden, 2022) to build custom code to analyze the peak evoked IPSC amplitude and produce trial-by-trial and averaged IPSC traces (available on GitHub). For each cell, peak IPSC amplitudes from 3-10 consecutive sweeps were averaged, and cell averages for edited and control conditions were compared in GraphPad Prism 10.0 using an unpaired t-test with Welch’s correction for unequal variances.

#### VP evoked IPSC/EPSC analysis

Evoked IPSC and EPSC recordings were exported in Axon Binary File (.abf) format. As above, we used the pyABF package to build custom code to determine the average current amplitude for 20ms around peak IPSC/EPSC deflection and produce trial-by-trial and averaged IPSC traces (available on GitHub). For each cell, the 20ms average values for 10-50 consecutive sweeps were averaged and the absolute value of the ratio of IPSC:EPSC amplitudes was calculated. Average EPSC amplitude and average IPSC:EPSC ratio were compared in GraphPad Prism 10.0 using an unpaired t-test with Welch’s correction for unequal variances.

#### VTA evoked NMDAR/AMPAR Analysis

AMPA and NMDAR currents were analyzed using Igor Pro software (Wavemetrics) with the NeuroMatic plug-in (Rothman & Silver, 2018). AMPAR currents were quantified as the average within 1 ms of the peak response. NMDAR currents were quantified 50 ms after the peak outward current. Currents represent the average amplitude from 10-15 consecutive sweeps recorded at 20 second intervals. Statistical comparisons were done using an unpaired t-test with GraphPad software.

All recordings and analysis were performed by investigators blinded to the group.

### Pavlovian Cue/Reward Task

The Pavlovian Cue/Reward task was performed in a clear 6.75”x12.25”x9.875” (WxLxH) plexiglass enclosure with a top-down camera monitoring mouse activity. For two weeks prior to the first recording session, mice were habituated to a reversed 12h dark:12h light cycle. For 3 days prior to the first recording session, mice were acclimated to the chamber and to the fiber optic cable attachment for 1 hour/day. No photometry recordings were made during this time. Between animals, the enclosure was cleaned with Clidox and water. 12 hours prior to the first recording session, food was removed from the animals’ home cage.

Animals were subjected to 5 30-minute-long recording sessions over the course of a week. During sessions, a FED3 device (Matikainen-Ankney et al., 2021) delivered a 4000Hz, 200ms tone followed 1-3s later by a food reward pellet. The tone was coupled with a TTL pulse to Bonsai to allow for alignment of photometry recording data with the tone cue. The mouse was freely able to take the pellet at will; reward acquisition was also coupled with a TTL pulse. Following reward acquisition, a variable 6-12s interval preceded the tone cue for the next trial. Mice were not pre-trained on this paradigm, and learned it over the course of the 5 trial days. Between trial days, animals received food *ad libitum* until 12h prior to the next day’s trial, at which point food was removed from their home cage.

### Fiber Photometry Setup & Analysis

Fiber photometry signals were collected using a 2m Doric fiber optic cable (Doric #D20714022). A Plexon 473nm LED Driver was used to deliver a constant 2.7 W blue light stimulus via the cable, and GCaMP8f photons were collected by the same fiber optic, passed through a dichroic mirror, and recorded in Bonsai. No isobestic control was used. Video was captured using a FLIR Camera (Sony IMX273, 226 FPS). Photometry signal, video, mouse (X,Y) position, and tone and reward TTL signals were recorded and synchronized using Bonsai. The photometry signal data was output as a .csv and analyzed using Guided Photometry Analysis in Python (GuPPy) (Venus N Sherathiya et al., 2021). Since no isobestic control was used, GuPPy used a moving average to calculate a baseline fluorescence curve, accounting for photobleaching during the recording. To analyze the average GCaMP8f fluorescence response to cue and reward stimuli, cue and reward times were extracted from the raw data and input into GuPPy. The raw fluorescence signal was normalized by dividing the change in fluorescence by the baseline fluorescence (ΔF/F), and the epoch from 2 seconds before to 2 seconds after each tone cue or reward acquisition was analyzed to determine the peak GCaMP8f response to cue or reward, respectively. An example input parameters document is available on GitHub. Heatmaps were generated in Python by averaging the GCaMP8f signal from 2 seconds before to 2 seconds after the cue or reward from all trials on all days for each subject. Overall cue and reward GCaMP8f responses were compared in Prism, where columns represented Control vs. Edited groups, and sub-columns represented each subject. The peak ΔF/F response to cue or reward for all trials was entered for each subject, and the Control vs. Edited responses were compared using an unpaired, 2-tailed nested t-test.

All photometry recordings and initial analysis in GuPPy were performed by investigators blinded to the group.

### Immunohistochemistry

#### VGAT

For VGAT immunohistochemistry in the NAc, mice were perfused with 40 mL chilled 1X PBS followed by 40 mL chilled 4% PFA. Brains were dissected out and post-fixed in 4% PFA overnight at 4°C. A Precisionary VF 310-0Z compresstome was used to cut 40 µm-thick coronal sections around the region containing the NAc. Sections were washed 3 times in 1X PBS, then incubated for 1 hour in blocking solution (0.3% TritonX (Sigma #X-100), 5% normal donkey serum (NDS). During incubation, a 1:1000 solution of mouse anti-VGAT antibody (SySy #131011) in blocking solution was made. Slices were incubated in VGAT antibody solution overnight at 4°C. The next day, slices were washed 3 times in 1X PBS, then incubated for 2 hours at 4°C in a 1:1000 solution of AF-568 donkey anti-mouse secondary antibody (Invitrogen #A10037) in blocking solution. Slices were washed 3 times in 1X PBS, then mounted on a slide and counter-stained with DAPI (SouthernBiotech #010020).

### *In situ* hybridization

#### VTA

To verify injection site, fiber placement, and GCaMP8f/Th overlap in VTA sections, we performed *in situ* hybridization (ISH) using the RNAScope Multiplex Fluorescence v2.0 kit (ACD #323110). Mice were perfused with 40 mL chilled 1X PBS followed by 40 mL chilled 4% PFA. Brains were dissected out and post-fixed in 4% PFA overnight at 4°C, then transferred to 30% sucrose in 1X PBS for overnight cryopreservation. Cryo-preserved brains were frozen in OCT (Sakura #4583) at -80°C, and 20 µm sections were taken on a Leica CM1860 cryostat. Sections were immediately mounted on glass slides, dried at room temperature for 1 hour, then stored at -80°C until processing. ISH was performed according to the RNAscope Multiplex Fluorescent Reagent Kit v2 User Manual, following tissue pretreatment instructions for fixed-frozen tissue. To identify cells expressing GCaMP8f mRNA from the virus injection, we used the EGFP-O4 Probe in Channel 1 (ACD #538851). To label *Th-*expressing cells in the VTA, we used the *Th* Probe in Channel 2 (ACD # 317621-C2). We used Opal 520 Reagent (Akoya #OP001001) and Opal 650 Reagent (Akoya #OP001005) to visualize the amplified EGFP-O4 and Th transcripts, respectively. After ISH, slices were coverslipped and with DAPI mounting media.

#### DRG

For analysis of infection efficiency, DRG sections were dried and ISH was performed using RNAscope, as above, with a probe against *Pvalb* (ACD #421931-C2) to identify target DRG neurons, and a probe against *mCherry* (ACD #431201) to identify infected neurons. The *Pvalb* probe was tagged with Opal 650 reagent (Akoya #OP001005) and the *mCherry* probe was tagged with Opal 570 reagent (Akoya #OP001003).

### Imaging

#### NAc imaging for VGAT puncta quantification

Experimenters were blinded to subject ID and condition during both imaging and analysis. Stained NAc sections were imaged on a Keyence BZ-X810 microscope at 60X magnification and high resolution with optical sectioning using a 10 µm 1D slit. DAPI was imaged with blue dichroic filter (Chroma #49021) at 1/5s exposure; GFP was imaged with a green dichroic filter (Chroma #49011) at 1/10s exposure, and VGAT (AF568) was imaged with a red dichroic filter (Chroma #49008) at 1/3.5s exposure for all sections across experiments with the same excitation intensity. All images were taken in areas with complete GFP expression to ensure accurate comparison of VGAT puncta counts between images.

#### VTA Imaging for virus expression and probe alignment

Stained VTA sections were imaged on a Keyence BZ-X810 microscope at 4X magnification. DAPI was imaged with blue dichroic filter (Chroma #49021) at 1/2.5s exposure; Opal 520 (GCaMP8f) was imaged with a green dichroic filter (Chroma #49011) at 1/3s exposure, and Opal 650 (*Th*) was imaged with a far-red dichroic filter (Chroma #49006) at 3s exposure. Mice with poor GCaMP8f expression were excluded from analysis. Mice where fiber placement was >100µm dorsal or lateral to GCaMP8f expression, and mice where fiber track was not located in the same slice as GCaMP8f expression, were also excluded from analysis.

#### Spinal Cord

PV-Cre/Cas9 mice were perfused at p14 using 1X PBS followed by 4% PFA, as above. Following perfusion, animals were decapitated and post-fixed in 4% PFA at 4°C overnight. Lumbar spinal cord and DRG were dissected out and post-fixed overnight, then cryopreserved in 30% sucrose at 4°C. Spinal cords and DRG were frozen separately in OCT at -80°C; 40 µm sections of both spinal cord and DRG were taken using a Leica CM1860 cryostat at -20°C and mounted on separate glass slides. Spinal cord sections were counter-stained with DAPI and imaged on a Keyence BZ-X810 microscope at 20X magnification, using optical sectioning with a 10 µm 1D slit. DAPI was imaged with blue dichroic filter (Chroma #49021) at 1/10s exposure and mCherry was imaged with a red dichroic filter (Chroma #49008) at 1/10s exposure. Images were stitched using BZ-X800 Analyzer, and stitched images were imported into FIJI for analysis.

#### DRG

DRG were imaged using a Keyence BZ-X810 microscope at high resolution and 40X magnification, using optical sectioning with a 10 µm 1D slit. DAPI was imaged with a blue dichroic filter (Chroma #49021) at 1/10s exposure, Opal 570 (*mCherry*) was imaged with a red dichroic filter (Chroma #49304) at 1/3.5s exposure, and Opal 650 (*Pvalb*) was imaged with a far red dichroic filter (Chroma #49006) at 1/1.2s exposure.

### Image Analysis

#### VGAT Puncta

Red channel images were uploaded into FIJI (FIJI is Just ImageJ) and analyzed as follows: first, pseudo-colored images were converted to 16-bit grayscale images. We then performed background subtraction with a rolling ball radius of 50 pixels. Images were converted to binary, using the threshold command such that all pixels with a value ≥9 were set to the maximum pixel value, and all values <9 were set to 0. Puncta were counted using the “Analyze Particles” command, selecting for puncta between 0.001-0.1 in^2^ and circularity between 0-1. The number of puncta per image was compared between control and edited groups using an unpaired, 2-tailed t-test with Welch’s correction for unequal variances in GraphPad Prism 10.0.

#### Spinal Cord

For analysis of mCherry+ projections to the spinal cord dorsal horn, stitched red channel images were converted to 16-bit black & white images, and background subtraction was performed with a rolling ball radius of a rolling ball radius of 50 pixels. ROIs were then drawn bilaterally around the point of fiber entry in the ventral horn, the intermediate zone, and the dorsal horn. We then calculated fluorescence intensity for each ROI and normalized it to the area in µm^2^. Images were then binarized with a threshold of 4, and the binary images were used to calculate the % area of mCherry+ fibers in each region. Fluorescence density and % Area values were averaged for each region in each subject and compared using an unpaired, two-tailed t-test with Welch’s correction for unequal variances.

#### DRG

Red and far-red channel images (representing *mCherry*+ and *Pvalb*+ cell bodies, respectively) were loaded into FIJI. Images were converted to 16 bit and background-subtracted using a 50 pixel rolling ball radius. *mCherry+* and *Pvalb+* cell bodies were identified manually, and the number of *mCherry+* cell bodies was divided by the number of *Pvalb+* cell bodies to obtain the fraction of *Pvalb+* neurons infected with virus. The fraction of *mCherry+/Pvalb+* neurons in each section was averaged for each subject, and average control and edited infection efficiency was compared in GraphPad Prism 10.0 using an unpaired, two-tailed t-test with Welch’s correction for unequal variances.

All tissue processing, imaging, and initial analysis in FIJI was performed by experimenters blinded to the group.

### Statistics

Experimenters were blinded during data collection and processing. All data were analyzed in GraphPad Prism 10.0. For all electrophysiology and imaging experiments, control and edited groups were compared using an unpaired, two-tailed t-test with Welch’s correction for unequal variances. Control and edited groups in fiber photometry experiments were compared using an unpaired, 2-tailed, nested t-test, where each sub-column represented one animal, and each data point within the sub-column represented the average of all trials for that animal on each recording day. All analysis code and GuPPy input parameters are available at https://github.com/jamiecm17/CRISPR-Cas9-Paper. All data are reported as mean±SEM.

